# Induction of C_4_ genes evolved through changes in *cis* allowing integration into ancestral C_3_ gene regulatory networks

**DOI:** 10.1101/2020.07.03.186395

**Authors:** Pallavi Singh, Sean R. Stevenson, Ivan Reyna-Llorens, Gregory Reeves, Tina B. Schreier, Julian M. Hibberd

**Author notes:** Both authors contributed equally (Pallavi Singh, Sean R. Stevenson).

## Abstract

C_4_ photosynthesis has evolved independently in over sixty lineages and in so doing repurposed existing enzymes to drive a carbon pump that limits the RuBisCO oxygenation reaction. In all cases, gene expression is modified such that C_4_ proteins accumulate to levels matching those of the photosynthetic apparatus. To better understand this rewiring of gene expression we undertook RNA- and DNaseI-SEQ on de-etiolating seedlings of C_4_ *Gynandropsis gynandra*, which is sister to C_3_ Arabidopsis. Changes in chloroplast ultrastructure and C_4_ gene expression were coordinated and rapid. C_3_ photosynthesis and C_4_ genes showed similar induction patterns, but C_4_ genes from *G. gynandra* were more strongly induced than orthologs from Arabidopsis. A gene regulatory network predicted transcription factors operating at the top of the de-etiolation network, including those responding to light, act upstream of C_4_ genes. Light responsive elements, especially G-, E- and GT-boxes were over-represented in accessible chromatin around C_4_ genes. Moreover, *in vivo* binding of many G-, E- and GT-boxes was detected. Overall, the data support a model in which rapid and robust C_4_ gene expression following light exposure is generated through modifications in *cis* to allow integration into high-level transcriptional networks including those underpinned by conserved light responsive elements.

## INTRODUCTION

Photosynthesis fuels most life on Earth and in the majority of land plants Ribulose 1,5-bisphosphate Carboxylase Oxygenase (RuBisCO) catalyses the initial fixation of atmospheric carbon dioxide (CO_2_) to generate phosphoglyceric acid. However, oxygen (O_2_) can competitively bind to the RuBisCO active site to form a toxic product 2-phosphoglycolate (Bowes, Ogren, and Hageman 1971). Although 2-phosphoglycolate can be metabolised by photorespiration (Bauwe, Hagemann, and Fernie 2010; Tolbert and Essner 1981) this takes place at the expense of carbon and energy and it is thought that the evolution of carbon concentrating mechanisms allowed rates of photorespiration to be reduced. C_4_ photosynthesis is one such example and is characterised by compartmentation of photosynthesis, typically between mesophyll and bundle sheath cells (Hatch 1987). This compartmentalisation involves cell-preferential gene expression and allows increased concentrations of CO_2_ to be supplied to RuBisCO sequestered in bundle sheath cells (Furbank 2011). Rather than RuBisCO initially fixing carbon, in C_4_ species fixation is initiated by phospho*enol*pyruvate carboxylase (PEPC) combining HCO_3_^-^ to form a C_4_ acid in the mesophyll. Diffusion of C_4_ acids into bundle sheath cells and subsequent decarboxylation results in elevated partial pressures of CO_2_ around RuBisCO facilitating efficient carboxylation and reducing the requirement for significant rates of photorespiration. C_4_ photosynthesis results in higher water and nitrogen use efficiencies compared with the C_3_ state, particularly in dry and hot climates. C_4_ crops of major economic importance include maize (*Zea mays*), sugarcane (*Saccharum officinarum*), sorghum (*Sorghum bicolor*), pearl millet (*Pennisetum glaucum*) and finger millet (*Setaria italica*) (Sage and Zhu 2011). Although C_4_ photosynthesis is a complex trait characterized by changes in anatomy, biochemistry and gene expression (Hatch 1987) it has evolved convergently from C_3_ ancestors in more than sixty independent lineages that together account for ∼8,100 species (Sage 2016). Parsimony would imply that gene networks underpinning this system are derived from those that operate in C_3_ ancestors.

Compared with the ancestral C_3_ state, expression of genes encoding components of the C_4_ pathway are up-regulated, and restricted more precisely to either mesophyll or bundle sheath cells. Our present understanding of these changes to C_4_ gene regulation is mostly based on studies designed to understand the regulation of individual C_4_ genes (Hibberd and Covshoff 2010). For example, a number of *cis*-regulatory motifs controlling the cell preferential expression of C_4_ genes have been identified (Brown et al. 2011; Burnell, Suzuki, and Sugiyama 1990; Glackin and Grula 1990; Gowik et al. 2004; Kajala et al. 2012; Reyna-Llorens et al. 2018; Williams et al. 2016). Whilst some *cis*-elements appear to have evolved *de novo* in C_4_ genes to pattern their expression (Matsuoka et al. 1994; Nomura et al. 2000, 2005; Stockhaus et al. 1997), others appear to have been recruited from pre-existing elements present in C_3_ orthologs (Brown et al. 2011; Kajala et al. 2012; Reyna-Llorens et al. 2018; Williams et al. 2016) and in these cases, there is evidence that individual *cis*-elements are shared between multiple C_4_ genes. In contrast to the analysis of regulators of cell specific expression, there is much less work on mechanisms that underpin the up-regulation and response to light of genes important for the C_4_ pathway compared with the ancestral C_3_ state. For example, whilst many C_4_ pathway components and their orthologs in C_3_ species show light-dependant induction (Burgess et al. 2016) the mechanisms driving these patterns are still largely unknown. One possibility is that *cis*-elements referred to as Light Responsive Elements (LREs) that have been characterized in photosynthesis genes in C_3_ plants (Acevedo-Hernández, León, and Herrera-Estrella 2005; Argüello-Astorga and Herrera-Estrella 1996) are acquired by genes of the core C_4_ pathway. The response of a seedling to light is the first major step towards photosynthetic maturity, but how C_4_ genes are integrated into this process is poorly understood. Seedlings exposed to prolonged darkness develop etioplasts in place of chloroplasts (Pribil, Labs, and Leister 2014). Etioplasts lack chlorophyll but contain membranes composed of a paracrystalline lipid–pigment–protein structure known as the prolamellar body (Bahl, Francke, and Monéger 1976). De-etiolation of seedlings therefore marks the initiation of photosynthesis and presents a good model system to study the dynamics and regulatory mechanisms governing photosynthesis in both C_3_ and C_4_ species.

Here, we used a genome-wide approach to better understand the patterns of transcript abundance and potential regulatory mechanisms responsible for these behaviours underpinning C_4_ photosynthesis. We integrate chromatin accessibility, predicted transcription factor binding sites, as well as binding evidence obtained *in vivo* and expression data to show that C_4_ pathway components in *Gynandropsis gynandra*, the most closely related C_4_ species to *Arabidopsis thaliana*, are connected to the top of the de-etiolation regulatory network. Photomorphogenic and chloroplast-related transcription factors were up-regulated upon exposure to light and showed highly conserved dynamics compared with orthologs from C_3_ Arabidopsis. Further analysis using an analogous dataset from Arabidopsis allowed us to compare the extent to which regulatory mechanisms are shared between the ancestral C_3_ and derived C_4_ systems. Firstly, this showed that the cistrome of C_4_ genes from *G. gynandra* has diverged from C_3_ orthologs in Arabidopsis. The C_4_ cistrome of *G. gynandra* showed an increased importance for homeodomain and Teosinte branched1/Cincinnata/Proliferating cell factor (TCP) transcription factors that may relate to the patterning of gene expression between mesophyll and bundle sheath cells in C_4_ leaves. However, the same C_4_ genes also acquired many LREs already associated with the regulation of C_3_ photosynthesis genes. Construction of a gene regulatory network recovered transcription factors from homeodomain, TCP as well as many LRE-related families as core components at the top of the de-etiolation network, and many were modelled to regulate C_4_ components. Together these data are consistent with C_4_ genes being rewired to allow their integration into existing gene regulatory networks that allow rapid, robust and spatially precise induction during de-etiolation.

## RESULTS

### De-etiolation and chloroplast development in C_4_ *Gynandropsis gynandra*

As etiolated seedlings of *G. gynandra* were transferred from dark-to-light, the dynamics associated with unfolding of the apical hook, chlorophyll accumulation, and ultrastructural re-arrangements of chloroplasts were determined. Classical photomorphogenic responses of apical hook unfolding and greening of cotyledons were visible 2 hours after transfer to light (Figure 1A). Chlorophyll accumulation was detectable by 0.5 hours after exposure to light, and that an initial exponential phase was followed by a more linear increase (Figure 1B). Little additional chlorophyll was synthesised from 12 to 24 hours after first exposure to light (Figure 1B). Assembly of the photosynthetic membranes in chloroplasts from mesophyll and bundle sheath cells, both of which are involved in C_4_ photosynthesis, was apparent over this time course (Figure 1C-D). In the dark, prolamellar bodies dominated the internal space of chloroplasts in each cell type. After 0.5 hours of exposure to light, although prolamellar bodies were still evident, they had started to disperse. Starch grains were apparent in bundle sheath chloroplasts by 24 hours after exposure to light, and it was noticeable that thylakoids showed low stacking in this cell type (Figure 1C-D, Supplemental Figure 1). Overall, these data indicate that in *G. gynandra*, assembly of the photosynthetic apparatus was initiated within 0.5 hours of exposure to light and by 24 hours the apparatus appeared fully functional. To better understand the patterns of gene expression and the transcriptional regulation that underpin this induction of C_4_ photosynthesis, these early time points were selected for detailed molecular analysis.

**Figure 1:**
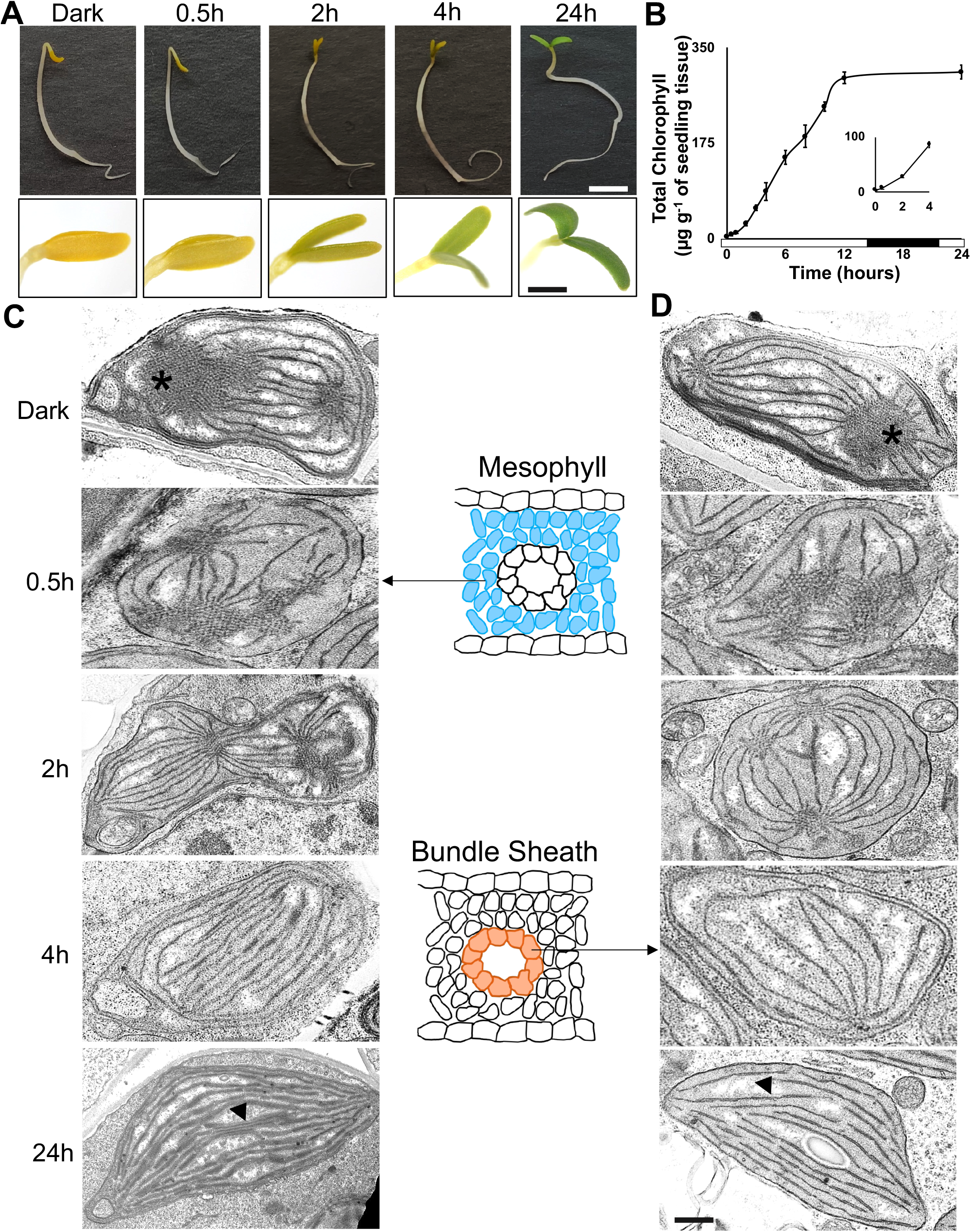
Establishment of photosynthesis in *G. gynandra*. **(A)** Representative images of *Gynandropsis gynandra* seedlings illustrating greening and unhooking of the cotyledons. **(B)** Total chlorophyll over the time-course (data shown as means from three biological replicates at each time point, ± one standard deviation from the mean). The first four hours show an exponential increase (inset). Bar along the x-axis indicates periods of light (0-14 hours), dark (14-22 hours) and light (22-24 hours). **(C-D)** Representative transmission electron microscope images of Mesophyll **(C)** and Bundle Sheath **(D)** chloroplasts of de-etiolating *G. gynandra* seedlings at 0, 0.5, 2, 4 and 24 hours after exposure to light. Asterisks and arrowheads indicate the prolamellar body and photosynthetic membranes respectively. Samples at each time were taken for RNA-SEQ and DNaseI-SEQ. Scale bars represent 0.5 mm for seedlings and 50 µm for cotyledons **(A)** or 500 nm **(C-D)**.

### Induction of photosynthesis genes in *G. gynandra*

To investigate how transcript abundance changed during the induction of C_4_ photosynthesis, mRNA from three biological replicates at 0, 0.5, 2, 4, and 24 hours after exposure to light was isolated and used for RNA-SEQ. On average, 10 million reads were obtained and ∼25,000 transcripts detected per sample (Supplemental Table 1). We were primarily interested in the dynamics of gene expression throughout de-etiolation and so analysed how transcript abundance changed relative to each previous time point. To provide a conservative estimate for the number of transcripts that were differentially expressed between consecutive time points, two independent algorithms were used and the intersect between these datasets determined (Supplemental Table 1). The greatest difference in transcript abundance was detected 0.5 hours after transfer from dark-to-light (number differentially expressed = 4609) with subsequent time points showing less than half this number (Supplemental Table 1). Principle component analysis (PCA) indicated replicates from each timepoint clustered together tightly, and that 64% of the variance in transcript abundance could be explained by two main components. The first accounted for 46% of the variance and was primarily associated with the dark-to-light transition, whilst the second accounted for 18% and was linked to time after transfer to light (Figure 2A). To provide insight into general patterns of differentially expressed genes (Supplemental Table 1), Gene Ontology (GO) terms were assessed (Figure 2B, Supplemental Figure 2, FDR < 10^−5^). Compared with each previous time point, up-regulated genes in samples taken at 0.5 and 24 hours showed enrichment in GO terms including those related to the plastid, as well as carbohydrate, secondary, nitrogen and lipid metabolism, but also responses to light and photosynthesis. These components were also over-represented in genes down-regulated at 2 and 4 hour time points largely due to most of the down-regulated genes at 2 hours previously having been up-regulated at 0.5 hours. This suggests that most early responding genes showed a dramatic change in transcription by 0.5 hours after which levels returned closer to pre-light levels. Presumably, this generates a large pool of mRNA templates for protein synthesis to allow assembly of the photosynthetic machinery.

**Figure 2:**
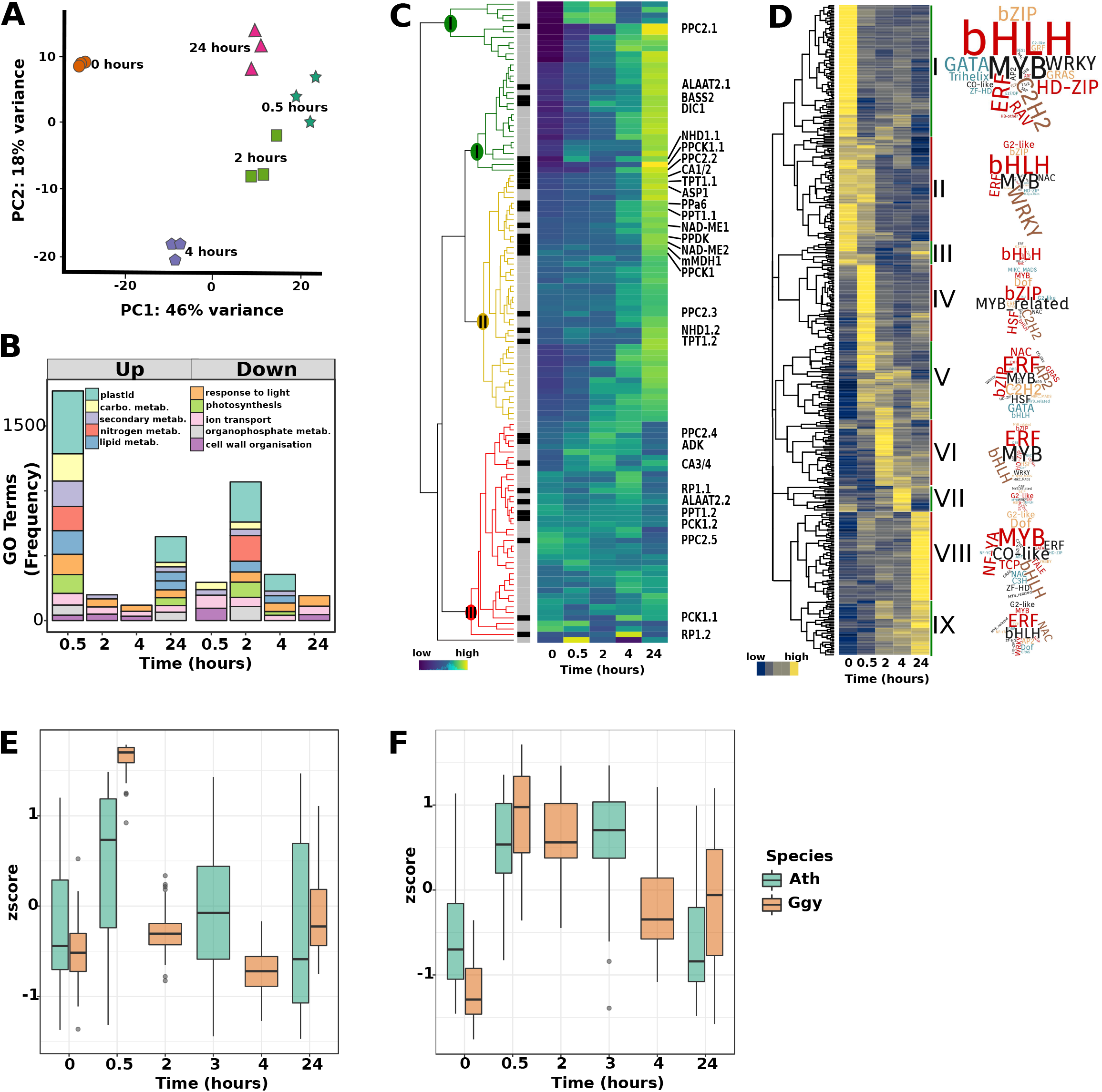
Changes in transcript abundance during greening of G. gynandra. **(A)** Principal component analysis of RNA-SEQ datasets. The three biological replicates from each timepoint of de-etiolating *G. gynandra* seedlings (0, 0.5, 2, 4 and 24 hours) formed distinct clusters. **(B)** Enriched GO terms between consecutive timepoints for up- and down-regulated genes. **(C)** Heatmap illustrating changes in transcript abundance of photosynthesis (grey sidebar) and C_4_ photosynthesis genes (black sidebar) during the time-course. Data are shown after normalisation of expression data with each gene plotted on a row and centred around the row mean. Colour-coding of the dendrograms (green, yellow and red) highlight expression clusters representing strong, moderate and no induction respectively. **(D)** Heatmap showing nine clusters containing all 433 differentially expressed transcription factors (TF). Values are shown as zscores with yellow and blue indicating higher and lower values respectively. Transcription factor families associated with each cluster are depicted in word clouds. **(E&F)** Transcription factors in clusters IV and V representing early induced regulators were compared to homologs in Arabidopsis. Expression values taken from Sullivan et al (2015) were plotted alongside *G. gynandra* data from this study. Note that time-points 0, 0.5 and 24 hours are shared but that 2 and 4 hours are unique to this study while 3 hours is unique to the Arabidopsis study. Expression zscores are shown for cluster IV **(E)** and V **(F)**.

To better understand the dynamics of gene expression associated with the induction of chlorophyll accumulation, remodelling of chloroplast ultrastructure in the C_4_ leaf and the establishment of C_4_ photosynthesis itself, genes associated with photosynthesis were subjected to hierarchical clustering. The genes defined as such were C_4_ pathway genes as well as orthologous nuclear genes from Arabidopsis annotated with the photosynthesis-related GO term (GO:0015979). A total of 116 genes were clustered into three main groups (Figure 2C) with clusters I and II showing strong and moderate induction respectively. Cluster III (red, Figure 2C) showed no clear induction over the time-course and whilst it did contain some putative C_4_ genes these represented poorly expressed paralogs of genes that were strongly induced and present in Cluster I or II. Re-analysis of an analogous de-etiolation dataset (Sullivan *et al*, 2014) from Arabidopsis showed that a smaller proportion of C_4_ genes responded to light (Supplemental Figure 3). Overall, these data show that C_4_ genes in both *G. gynandra* and Arabidopsis were induced after exposure to light but in *G. gynandra* a greater number responded strongly.

**Figure 3:**
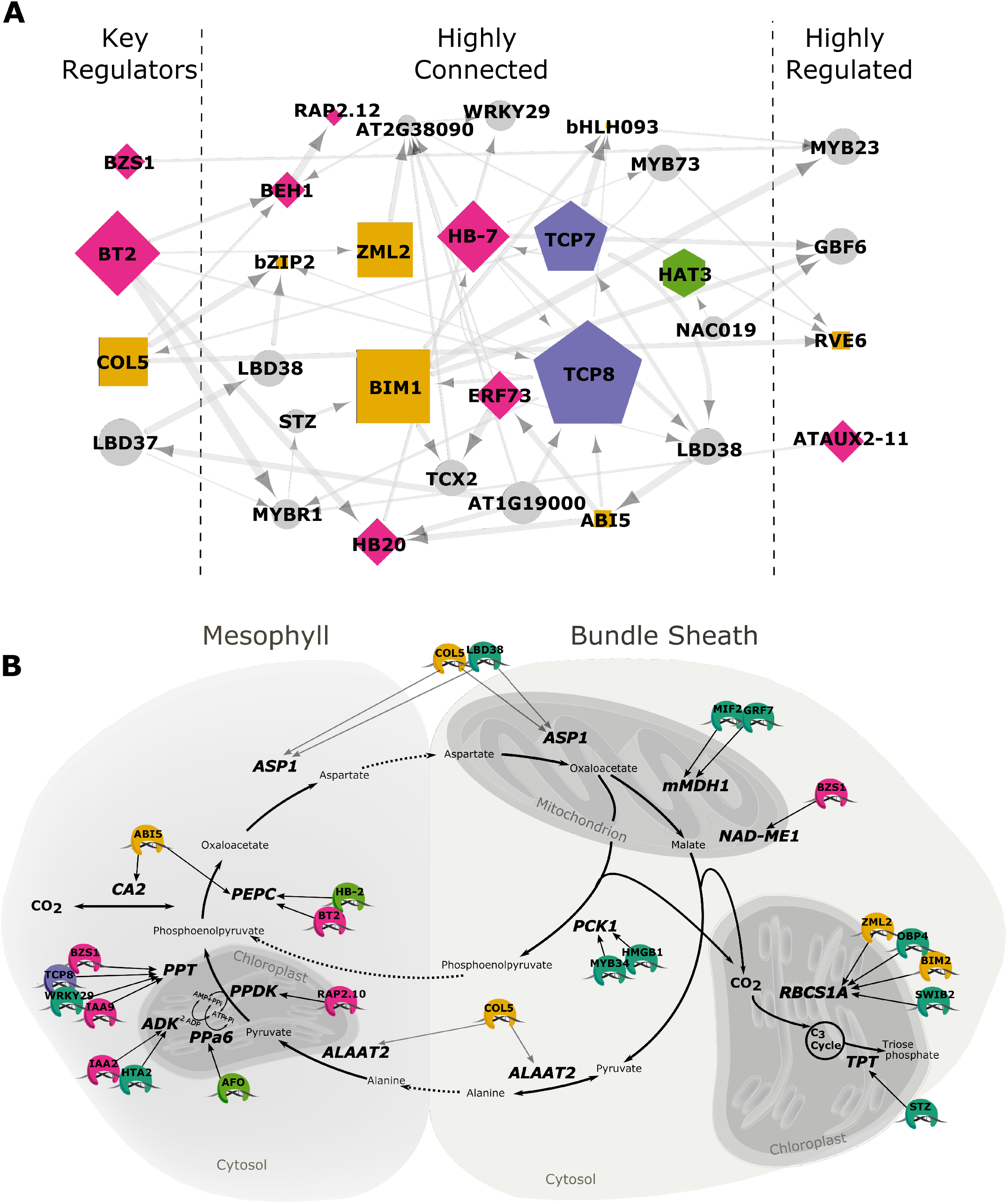
Time-series network construction of *G. gynandra* transcription factors and C_4_ pathway components. **(A)** Transcription factor network built from the most dynamic transcription factors across the 0, 0.5, 2 and 4 hours time-points. Networks were built using the dynGENIE3 and SWING algorithms with the edge weights combined to create an ensemble network. Transcription factors presented were with those the most in- and out-edges as well as highest betweenness centrality (BC) scores. Arrows indicate direction of interaction with regulators and predicted targets with thicker lines indicating higher edge weights and larger nodes indicate higher connectivity. Gene names shown are based on closest Arabidopsis homologs with the following broad functional groupings: Yellow squares; light regulation, green hexagons; developmental and patterning, purple pentagons; TCP, pink diamonds; phytohormone signalling, grey circles; other. **(B)** Transcription factors predicted to be upstream of each C_4_ gene. Genes encoding proteins of the C_4_ cycle are shown. Names and colours of transcription factors are defined in the same manner as **(A)**.

To better understand the potential mechanisms of transcriptional regulation during de-etiolation, we clustered 434 highly dynamic transcription factors (Figure 2D) that were differentially expressed between each consecutive time-point (Supplemental Table 2). This resulted in nine clusters with differing peaks in expression across the time-course. Clusters I to III represented around a third of the transcription factors, were down-regulated in response to light and contained many bHLHs transcription factors (Figure 2D). BIRD/INDETERMINATE DOMAIN and GAI-RGA-and-SCR (GRAS) transcription factors thought to play roles in regulating cellular leaf patterning, as well as many Homeodomain (HD)-Zip and Zinc Finger Homeodomain (ZFHD) factors were also down-regulated in response to light. In contrast, cluster IV peaked after 0.5 hours of light and included many bZIP, HSF and MYB-related transcription factors from the CCA1 family. Cluster V contained genes induced at 0.5 hours but expression was more sustained. Notably, clusters IV and V included orthologs of well-known regulators of chloroplast development such as GNC and GLK2 (Zubo et al. 2018) as well as the master regulator of de-etiolation and photomorphogenesis HY5 (Gangappa and Botto 2016). Clusters VI through IX contained transcription factors that peaked at each subsequent time point, and so likely represent later acting transcription factors in the response to light. By 24 hours, initial events in building the photosynthetic apparatus have taken place (Figure 1) and so transcription factors in clusters VIII and IX are likely involved in maturation and maintenance.

We next sought to quantify the degree of conservation in expression patterns following the onset of light between C_4_ *G. gynandra* and C_3_ Arabidopsis. We focused on clusters IV and V as their expression and annotations were them representing early regulators of de-etiolation. *G. gynandra* and Arabidopsis transcription factors showed strong conservation in their expression patterns. Orthologs from cluster IV showed a clear peak at 0.5 hours that was not sustained (Figure 2E) while those from cluster V showed a longer lasting induction in both species (Figure 2F). These data are consistent with the notion that the behaviour of transcription factors acting in networks associated with de-etiolation are conserved between C_3_ Arabidopsis and C_4_ *G. gynandra*.

In summary, consistent with the synthesis of chlorophyll and maturation of photosynthetic membranes of the chloroplast, photosynthesis-related genes were rapidly induced during de-etiolation. Moreover, early responding transcription factors from *G. gynandra* showed similar dynamics to homologs in Arabidopsis and belonged to families known to govern de-etiolation. As C_4_ pathway genes showed similar induction dynamics to C_3_ photosynthesis genes and the transcription factor behaviours were conserved, it would appear most likely that C_4_ genes have acquired mutations in *cis* such that they become incorporated into the pre-existing gene regulatory networks associated with de-etiolation.

### The C_4_ cycle and transcription factors networks

We next sought to define a core transcription factor network associated with the earliest timepoints of *G. gynandra* de-etiolation, and then investigate whether components of this network act upstream of C_4_ genes. Two algorithms namely SWING (Finkle, Wu, and Bagheri 2018) and dynGENIE3 (Huynh-Thu and Geurts 2018) were used to build a time-series network of the transcription factors in *G. gynandra* using expression patterns during the first 4 hours (0, 0.5, 2 and 4 hours). The network is generated from predicted interactions based on regression modelling of transcript abundance of all pairs of transcription factors and in so doing assigns a likelihood score (edge-weight) for each factor regulating another. Such an approach allows for the incorporation of time-delays between effectors and target dynamics which is not possible in traditional Gene regulatory Networks (GRNs) built from correlation scores (Finkle et al. 2018; Huynh-Thu and Geurts 2018). We filtered the network to focus on components predicted to regulate many down-stream targets (many out-edges), those regulated by many other transcription factors (many in-edges), and also how “central” a factor was (high betweenness centrality, BC) (Figure 3A). Transcription factors predicted to regulate many down-stream targets included orthologs of BT2, COL5, LBD37 and BZS1 (left hand segment, Figure 3A) and those predicted to have many regulators included GBF6, MYB23, AUX2-11 and RVE6 (right hand segment, Figure 3A). The most highly connected transcription factors (many in- and out-edges) tended also to have the highest BC scores, and included bHLH BIM1, GATA ZML2, HB-7, TCP8, TCP7 and ERF73 (central segment, Figure 3A). Notably, the network contained a range of transcription factors implicated in the regulation of photosynthesis genes in response to light including G- and E-box binding bZIP, bHLH and BZR transcription factors, the GATA ZML2, the BBX COL5, and also the clock-related RVE6 (yellow squares, Figure 3A). We also found an ortholog to the homeobox factors HAT3 (green hexagon, Figure 3A) which is a regulator of the dorsal-ventral axis in both cotyledons and developing leaves (Bou-Torrent et al. 2012). Overlaying this were transcription factors with annotations associated with phytohormone pathways such as ABI5, HB20 and BT2 (ABA), ERF73 and RAP2.12 (ethylene), BEH1 and BZS1 (brassinosteroids) and AUX2-211 (auxin) (pink diamond, Figure 3A). All of these represent transcription factors that are likely to play important roles in orchestrating the expression of others during de-etiolation with some likely acting as hubs (*e*.*g*., large central nodes, Figure 3A) and early master regulators triggering the shift to de-etiolation.

We next sought to integrate the transcription network with C_4_ pathway genes to assess how they may be connected to these central regulators during de-etiolation. For each C_4_ gene we kept the highest scoring edge as well as any other high scoring edges above a threshold such that each C_4_ component had one or more of the best predicted transcriptional regulators (Figure 3B). These networks included many of the transcription factors recovered from the original network (Figure 3A) suggesting their expression patterns were predictive of C_4_ genes. A small number of transcription factors were not present in the original network but were predicted to be upstream of a C_4_ gene. This included orthologs to HB-2 and AFO upstream of *PEPC* and *PPa6* respectively that have been implicated in early leaf development (Siegfried et al. 1999; Turchi et al. 2013). Overall, these networks reflect a complex regulatory hierarchy with both direct and indirect connections as well as potential “feed-forward” motifs. Importantly, this analysis of the expression data does not suggest a clear “master regulator” for C_4_ components (Westhoff and Gowik 2010) but rather a collection of transcription factors many found within the central transcription factor network. While the proposed edges are not proof of direct regulation, they identify individual transcription factors that have the strongest predicted effect on induction of C_4_ pathway genes during de-etiolation. In summary, we identified a core transcription factor network that likely plays important roles during de-etiolation of C_4_ *G. gynandra*. Many of the transcription factors in this network relate to light, hormones and leaf development and some are also predicted to regulate C_4_ pathway genes.

### Chromatin dynamics associated with de-etiolation in *G. gynandra*

While expression modelling generated predictions about gene interactions and transcription factor networks, we sought to refine these findings by analysing potential transcription factor-DNA interactions. To do so we used genome wide DNaseI-SEQ to first define regions of chromatin available for transcription factor binding and second to test whether *cis*-elements bound by components of the gene regulatory network presented were over-represented in these regions of accessible chromatin. Nuclei from three biological replicates across the five time points, as well as de-proteinated DNA controls were isolated. We first focused on understanding patterns in chromatin accessibility and potential *cis*-elements within such regions. From these time-points, a total of 1,145,530,978 reads were mapped to the *G. gynandra* genome, and 795,017 DNaseI-hypersensitive sites (DHS) representing broad regulatory regions accessible to transcription factor binding were identified (Figure 4A, Supplemental Figure 4). The average length of these DHS was ∼610 base pairs, and distribution plots showed that their density was highest at the predicted transcription start sites (Figure 4B). However, consistent with the notion that exposure to light leads to a rapid increase in open chromatin around gene bodies (Liu et al. 2017), peak DHS density at transcription start sites more than doubled by two hours after transfer to light (Figure 4B). We calculated the proportion of overlap between normalised DHS at each time-point and found high levels of change between 0 and 0.5 hours and 0 and 2 hours (64% and 71% non-overlapping) but less between 4 and 24 hours (40% non-overlapping) (Figure 4C). This suggests that chromatin dynamics decreased after an initial period of volatility coincident with a transcriptional burst following light exposure. Despite these global patterns, changes in DHS position were poorly predictive of gene expression (Supplemental Figure 5) suggesting that remodelling of transcription factors binding within these regions is a stronger determinant of gene expression.

**Figure 4:**
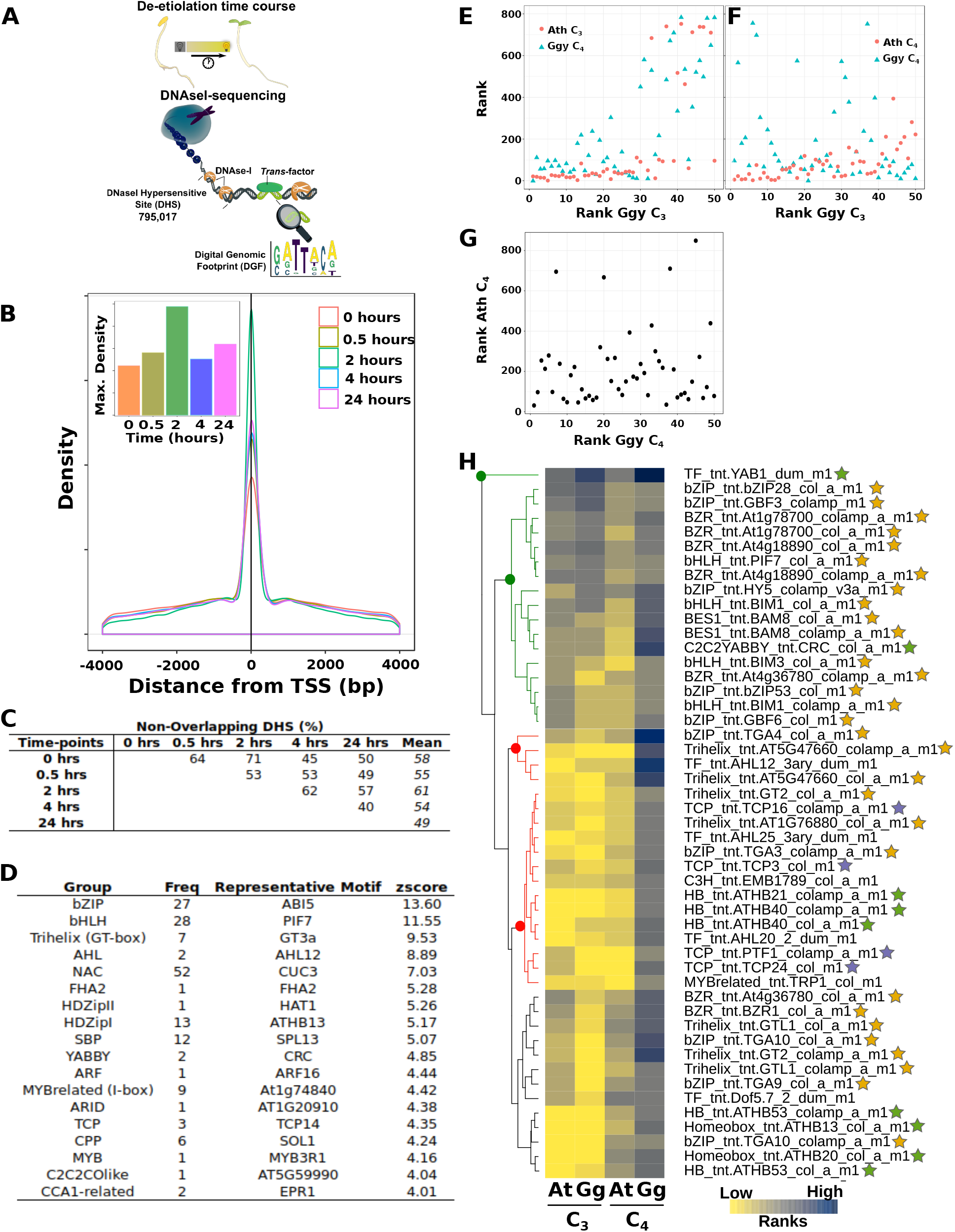
Profiling of open chromatin during de-etiolating of *G. gynandra*. **(A)** Schematic illustrating DNaseI-SEQ and total number of DNaseI-hypersensitive sites (DHSs) detected. **(B)** Density of open chromatin plotted relative to the nearest Transcription Start Site (TSS). Inset highlights maximum density overlapping with the TSS at each time point. **(C)** Percentage of non-overlapping DHSs at each timepoint. **(D)** Table showing the significantly enriched motifs in the DHS of significantly up-regulated genes at 0.5 hours. Groups represent similar motifs with the number of significantly enriched motifs in each group shown. A representative motif is shown with zscore (high values represent strong enrichment). bZIP, bHLH and Trihelix groups contain the most strongly enriched motifs. **(E-G)** Scatter plots showing the most enriched motifs in each cistrome (where 1 represents the most enriched motif). **(E)** Top 50 motifs in photosynthesis genes of C_3_ Arabidopsis (At) and C_4_ *G. gynandra* (Gg), **(F)** C_4_ and photosynthesis genes of C_3_ Arabidopsis, and **(G)** C_4_ genes from C_3_ Arabidopsis and *G. gynandra*. Motifs from the cistromes of C_3_ and C_4_ genes that showed induction during de-etiolation. **(H)** Heatmap of the top 50 motifs from DHS of C_4_ genes in *G. gynandra* compared with their ranking in C_4_ genes of Arabidopsis and photosynthesis genes in both species (log2 of the motif ranks across all four cistrome sets). Two distinct groups are highlighted with green motifs being highly ranked (more enriched) in all four cistromes while red motifs are those specifically highly ranked in the *G. gynandra* C_4_ cistrome. Broad functional groups are annotated tellow; light regulation (e.g., LRE), green; YABBY/homeodomain, purple; TCP.

**Figure 5:**
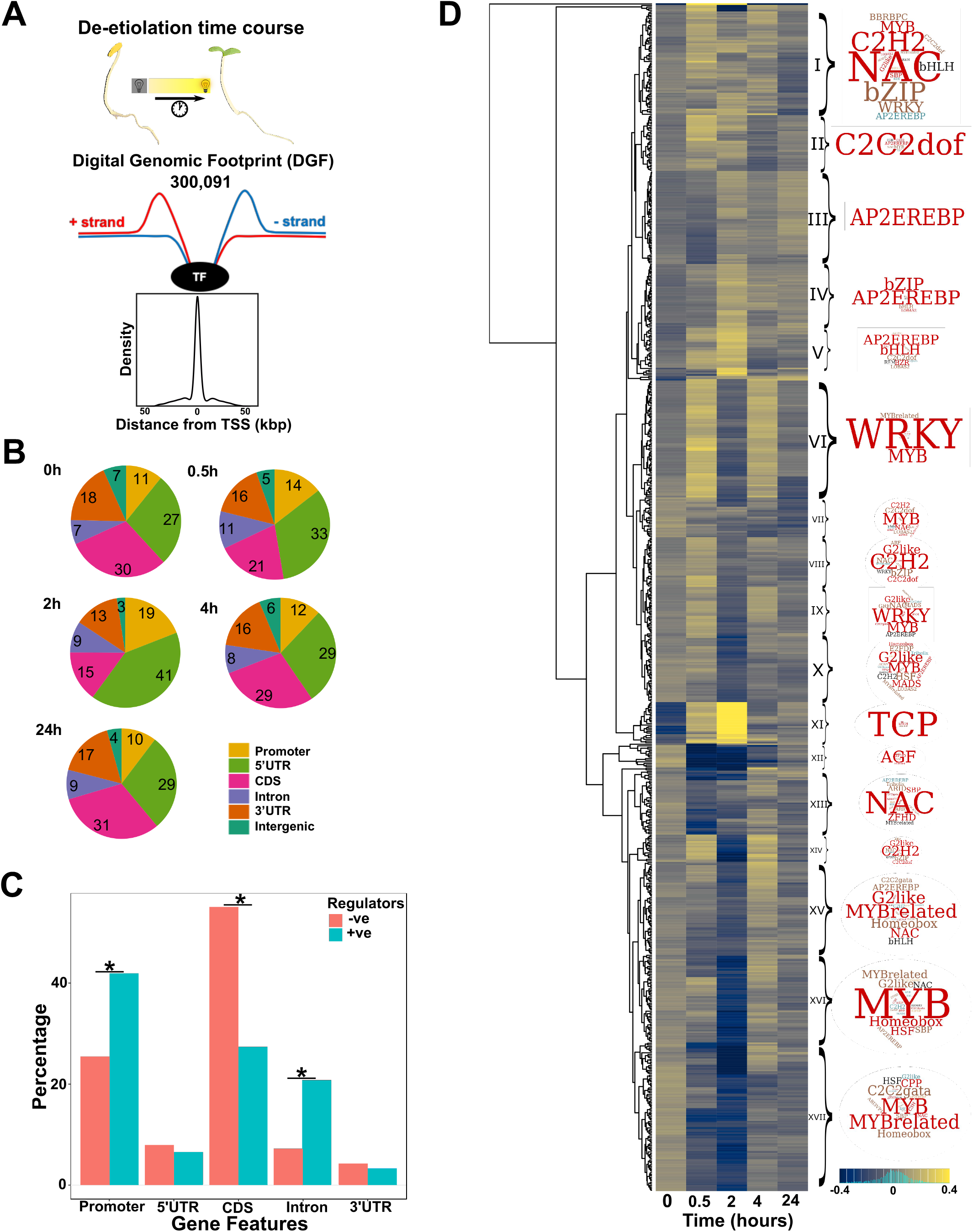
Transcription factor binding atlas for de-etiolating seedlings of *G. gynandra*. **(A)** Schematic illustrating sampling, number of Digital Genomic Footprints (DGF) identified and representative density plot of DGF positions relative to the nearest transcription start site (TSS). **(B)** Pie-charts summarising the density of DGF among genomic features. Promoters were defined as sequence < 2000 base pairs upstream of TSSs while intergenic represent any regions not overlapping with other features. Values indicate proportion of DGFs in each feature. **(C)** Bar chart showing the percentage of DGFs predicted to function either as activators (coral bars) or repressors (turquoise bars) lying within gene features of target genes. Statistically significant differences were found for promoters, CDS and intronic regions using a Chi square goodness of fit test (“*”). **(D)** Heatmap of motif frequencies (log10 sample normalised motif frequency/row mean) during de-etiolation. To illustrate identity and heterogeneity of motif groups, clusters were annotated with Wordclouds.

In order to identify potential regulatory mechanisms allowing genes to respond rapidly to light, the DHS of all significantly up- and down-regulated genes at 0.5, 2 and 4 hours were assessed for motif content. DHS of genes down-regulated after 0.5 hours of light included many AP2, WRKY, ZFHD and other homeobox, TCP, LOBAS2, BPC, TGA bZIP and GATA motifs (Supplemental Table 3). DHS of genes up-regulated after 0.5 hours of light were enriched in motifs associated with light, and included G-(bZIP), E-(bHLH and BZR), GT-(Trihelix) and I-box (MYB-related) motifs as well as CCA1-like (evening element) motifs (Figure 4D). Other motifs enriched at 0.5 hours included CPP, C2C2YABBY, homeobox and NAC motifs. To more clearly identify motifs most likely to be involved in the strong and rapid induction of gene expression in response to light we identified motifs enriched in 158 genes that were induced from 0 to 2 hours after exposure to light. This highlighted the role of G-boxes (bZIP motifs) as well as SBP, TCP and FAR1 motifs. While no motifs were statistically enriched in genes up regulated at both 2 and 4 hours, the TCP motifs were among the most common implicating a role for this group in the later stages of de-etiolation (Supplemental Table 3). In summary, the data are consistent with outputs of the gene regulatory network in which transcription factors downstream of the light response were predicted to be key regulators of early transcriptional responses during de-etiolation of C_4_ *G. gynandra*.

We next sought to understand whether photosynthesis genes could be linked to these canonical light responses, and particularly whether this was the case for genes of the C_4_ pathway. By re-analysing publicly available data for C_3_ Arabidopsis we aimed to investigate whether changes in *cis* are associated with the strong induction of C_4_ genes in *G. gynandra*. C*is*-elements in DHS associated with C_4_ orthologs (the C_4_ cistrome) from each species were identified. As well as genes encoding the core C_4_ pathway, we also included classical photosynthesis associated nuclear genes that showed clear induction in response to light (the C_3_ cistrome, Figure 2C). These gene sets allowed us to investigate the extent to which C_4_ genes in *G. gynandra* share a *cis*-code with photosynthesis genes in general, and whether this code is also found in the C_3_ ancestral state. As the number of any motif can vary between species due to phylogenetic distance, we ranked motif enrichment in each species. Of the fifty most enriched motifs from the *G. gynandra* C_3_ cistrome, the majority were also highly ranked in the Arabidopsis C_3_ cistrome (Figure 4E) indicating conservation of motifs associated with the regulation of photosynthesis in these species. This included many bZIP (including HY5), bHLH (including PIF7) and BZR motifs (Supplemental Table 4) supporting a model in which photosynthesis genes are direct targets of these early activators associated with de-etiolation. The cistromes of C_3_ and C_4_ genes from *G. gynandra* were less similar than the C_3_ cistromes of Arabidopsis and *G. gynandra* (Figure 4E) indicating that C_4_ genes from *G. gynandra* have not simply converged on the *cis*-code associated with C_3_ photosynthesis. Further, the *G. gynandra* C_4_ cistrome was less similar to the *G. gynandra* C_3_ cistrome than that of the Arabidopsis C_4_ orthologs (Figure 4F). Lastly, the *cis*-code of C_4_ genes from *G. gynandra* and Arabidopsis showed little conservation (Figure 4G) consistent with C_4_ genes being repurposed in the derived C_4_ state.

We next assessed annotations associated with the fifty most common motifs in the C_4_ cistrome of *G. gynandra* and linked these binding sites back to the expression data. Hierarchical clustering revealed two groups of particular interest (Figure 4H). The first (green; Figure 4H) contained motifs that were relatively highly ranked in all cistromes except C_4_ genes from Arabidopsis. This group included YAB1 and CRC, as well as bZIP G- and bHLH E-box motifs (Figure 4H). As all these motifs were enriched in DHS of genes up-regulated at 0.5 hours and the Arabidopsis C_4_ genes were more poorly expressed during de-etiolation, we suggest these motifs represent strong candidates for the early light induced expression of C_4_ genes from *G. gynandra* as well as C_3_ photosynthesis genes from both species. The second group (red) contained motifs that were only highly ranked in the C_4_ cistrome of *G. gynandra* and included several TCP and homeodomain as well as the GT-box binding trihelix and bZIP TGA motifs (Figure 4H). Most of these DNA binding sites were also enriched in rapidly induced genes in *G. gynandra*, with TCP motifs showing a stronger enrichment in those genes up regulated at two hours. The remaining motifs showed intermediate patterns and included additional G-box binding bZIP, GT-box binding trihelix, E-box binding BZR and a number of homeodomain motifs (Figure 4H).

In summary, the most enriched motifs in DHS of C_4_ pathway genes showed strong overlap with those found in genes rapidly induced during de-etiolation (Figure 4D). Furthermore, many transcription factors predicted to recognise these motifs were in early responding clusters (Figure 2D-F) and in the core transcription factor network (Figure 3A). Some motifs were enriched in *G. gynandra* C_4_ genes as well as C_3_ genes of both species while others were uniquely enriched in the C_4_ genes of *G. gynandra*. We conclude that C_4_ genes of *G. gynandra* are highly connected to the top of the de-etiolation regulatory network allowing rapid and strong expression during the earliest stages of de-etiolation, however many of these connections were not associated with C_3_ photosynthesis genes.

### A *cis-*regulatory atlas for de-etiolation in *G. gynandra*

Chromatin accessibility assays followed by *in silico* analysis of motifs in regions of open chromatin identified regulatory elements that could be important for gene regulation but cannot indicate whether motifs are actually subject to transcription factor binding. We therefore carried out sequencing at sufficient depth to define DNA sequences that are protected from DNaseI digestion (Figure 5A). Such sequences are diagnostic of strong and/or widespread protein binding and referred to as Digital Genomic Footprints (DGFs). Although DNaseI-SEQ has been used to predict transcription factor binding sites, the DNaseI enzyme possesses sequence bias and so to account for this de-proteinated DNA was analysed to enable footprint filtering (He et al. 2014; Yardimci et al. 2014) (Supplemental Figure 4). After filtering, 300,091 DGFs corresponding to individual transcription factor binding sites across all time points were identified (Figure 5A and Supplemental Figure 4).

The distribution of DGFs in gene features (*e*.*g*., promoters, exons, introns and UTRs) of *G. gynandra* changed during de-etiolation (Figure 5B). Notably, in the first 2 hours of exposure to light, DGF density in promoter elements (defined as sequence two kilobase pairs upstream of predicted transcriptional start sites) and 5′ UTRs increased (from 11 to 19% in the case of promoters and 27 to 41% for 5′ UTRs). This finding is consistent with the increase in DHS density around predicted transcriptional start sites at this time (Figure 2B). Coincident with the increase in DGFs in promoters and 5′ UTRs, the density found in coding sequence was reduced by around half (Figure 5B). In contrast, the density of DGFs associated with introns and 3′ UTRs changed little during de-etiolation. These findings suggest changes to binding site distribution between genomic features may play an important role in transcriptional regulation and contribute to the induction of photosynthesis during de-etiolation. To test this further, we correlated the change in frequency of each motif with change in expression of the nearest gene (Supplemental Table 5). Positive and negative correlations between motif frequency and gene expression were used to classify motifs as either allowing activation or repression. Of the motifs predicted to act as activators several Cysteine-rich polycomb-like protein (CPP) transcription factors were also found to be enriched in the 0.5 hour up-regulated DHS (Figure 4D), and significantly more were located in promoters and introns. In contrast, motifs predicted to act as repressors were around two-fold more likely to be found in exons (Figure 5C, Supplemental Table 5). Motifs with no correlation to targets were found to have intermediate frequencies suggesting a gradient between the two extremes (Supplemental Table 5).

To understand the dynamics associated with motif binding during de-etiolation, low frequency motifs were removed and the remaining 743 subjected to hierarchical clustering (Figure 5D). Although a few clusters were dominated by a small number of motifs bound by a particular family of transcription factor, most contained motifs bound by multiple families (Figure 5D). Cluster XI was made up of DGF predicted to be bound by TCP factors and showed a striking pattern of maximum abundance at 2 hours. This supports the finding that TCP motifs were enriched in genes up-regulated at 0.5 and more strongly at 2 hours. Hierarchical clustering split the DGF into two broad groups - those that increased in activity at 0.5 or 2 hours following light, and those that decreased in activity after exposure to light. Clusters I, IV and V showed an increase after light and were predicted to be bound by bZIP and bHLH transcription factors. These patterns in the DGF data support the motif enrichment analysis suggesting that not only do the accessible regions of up-regulated genes contain many bZIP and bHLH binding sites (G- and E-boxes respectively), but that many are actively bound. Deep DNaseI sequencing and foot-printing therefore supported patterns observed from the cistrome analysis but in addition allowed a direct read-out of transcription factor activity.

### Regulation of C_4_ genes by light responsive elements and their cognate transcription factors

By combining outputs from the RNA-SEQ and DNaseI-SEQ outlined above, we propose a model (Figure 6A) to explain the strong induction of C_4_ genes in *G. gynandra*. This is based on the observation that transcripts encoding certain families of transcription factors were strongly induced after 30 minutes of exposure to light, but also that motifs associated with these proteins were enriched in the cistrome of C_4_ genes, and also that in some cases *in vivo* binding by transcription factors was detected. By combining these data, we propose that C_4_ genes in *G. gynandra* become strongly induced early on in de-etiolation because they are wired such that they respond to a broad set of light regulatory factors including bHLHs, bZIPs and GT trihelixes (Figure 6A&B). Moreover, networks unrelated to canonical light signalling, but instead involving homeodomain and TCP transcription factors were also linked to the C_4_ genes (Figure 6A&B) and it seems likely that these additional motifs regulate aspects of the C_4_ pathway such as the patterning of gene expression to either mesophyll of bundle sheath cells.

**Figure 6:**
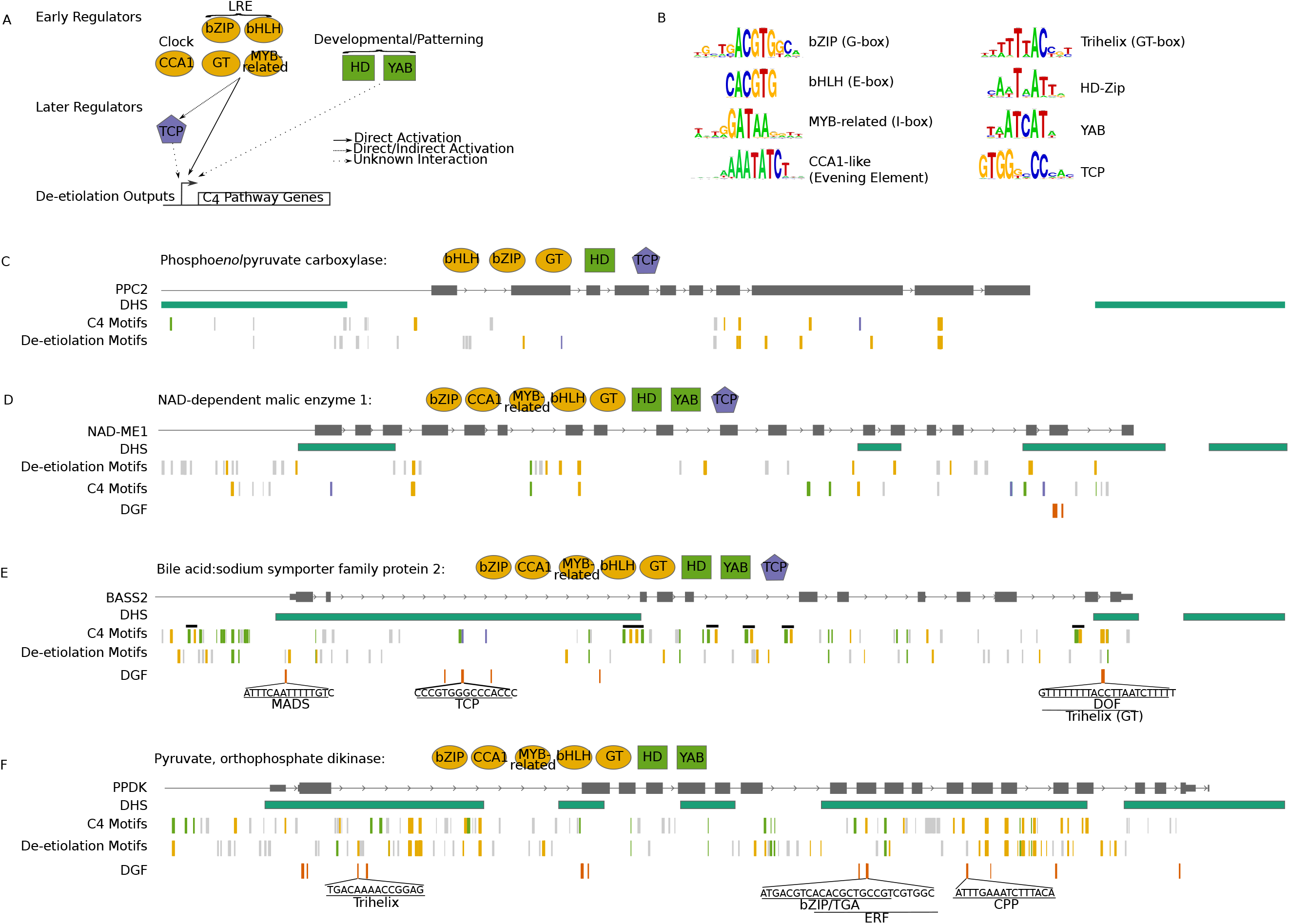
Model summarising proposed regulation of C_4_ genes. **(A)** Schematic of transcription factors predicted to regulate C_4_ pathway genes during de-etiolation. Transcription factor groups relating to light or clock regulation are shown in yellow, TCP in purple and developmental/patterning in green. LRE related transcription factors predicted to be activators may directly or indirectly activate TCPs. **(B)** Consensus motif sequence logos for each of the eight transcription factor groups. Schematics of PPC2 **(C)**, NAD-ME1 **(D)**, BASS2 **(E)** and PPDK **(F)** gene models with DHS regions, predicted enriched motifs and *in vivo* binding sites (DGF) annotated. Transcription factor groups predicted to bind to notable motifs are shown above schematics next to each gene name. C _4_ motifs relate to the most enriched motifs found in the DHS regions of all C_4_ pathway genes (see Figure 4H) while de-etiolation motifs relate to those found in the 0.5 hours up-regulated genes (see Figure 4D). *In vivo* evidence of binding is shown as DGF sites and their sequences are shown where they overlap enriched motifs.

The data also indicate that individual C_4_ genes differ in how, and the extent to which, they are plumbed into these gene regulatory networks. This conclusion is exemplified by four genes encoding core components of the cycle: *PEPC, NAD-ME1, BASS2* and *PPDK*. In the case of *PEPC*, which carries out the first committed step of the C_4_ pathway, two DHS in close proximity to the gene were sparsely populated with the motifs summarised above (Figure 6C). However, LRE binding sites were detected outside of the DHS (Figure 6C). NAD-ME1, a decarboxylase in the bundle sheath, was associated with a DHS overlaying the proximal promoter and the 5′ of the gene body as well as further highly accessible regions at the 3′ of the gene body (Figure 6D). Numerous LRE binding sites including G-, E-, I- and GT-boxes and a number of HD and YAB and TCP sites were detected. BASS2 that allows pyruvate uptake by chloroplasts in mesophyll cells was associated with a large DHS overlaying the predicted proximal promoter and 5′ of the gene (Figure 6E). The gene contained a high number of LRE, some of which were in the DHS, but others which were distributed across the gene body. BASS2 contains two TCP motifs, one of which was bound *in vivo*. It was also noticeable that many HD and GT-box sites were adjacent to one another suggesting these two motifs may act as a complex (black bars, Figure 6E). Lastly, PPDK which allows PEP to be regenerated from pyruvate was associated with a number of DHS, and these contained a relatively high density of LRE, three of which were bound *in vivo* (Figure 6F). PPDK therefore appears to be both highly accessible and richly endowed with LRE binding sites.

Overall, the data we present provide insights relating to the regulation of C_4_ genes, and how this has likely evolved from the ancestral C_3_ state. Many aspects of the existing *cis*-code associated with the response of C_3_ photosynthesis to de-etiolation appears to have been acquired by genes of the C_4_ pathway. In addition, C_4_ genes have also acquired motifs associated with transcription factors unrelated to light signalling or C_3_ photosynthesis genes, but rather linked to families of transcription factors implicated a wide range of functions. We propose that in both cases these changes in *cis* have allowed C_4_ genes to be rewired such that they become integrated with transcription factors at the top of existing de-etiolation gene regulatory networks. We therefore conclude that alterations in *cis* are fundamental in allowing C_4_ genes to be patterned to mesophyll and bundle sheath cells but at the same time co-regulated with genes encoding the photosynthetic apparatus.

## DISCUSSION

### C_4_ genes of *G. gynandra* are induced rapidly during de-etiolation

The dark-to-light transition has been frequently used to understand assembly of the photosynthetic apparatus (Armarego-Marriott et al. 2019; Dubreuil et al. 2018). As the cotyledons of *G. gynandra* operate C_4_ photosynthesis (Koteyeva et al. 2011), de-etiolation of this species appeared an attractive system with which to probe how C_4_ genes are co-regulated with other photosynthesis genes. Consistent with this, and with previous analyses of species that use the ancestral C_3_ pathway (Bahl et al. 1976; Baker and Butler 1976; Boardman 1977; Boffey, Selldén, and Leech 1980; Krupinska and Apel 1989), chloroplasts from *G. gynandra* followed a trajectory towards full photosynthetic capacity over twenty-four hours. The induction of photosynthesis transcripts in *G. gynandra* within 0.5 hours of exposure to light was associated with up-regulation of homologs of master regulators of de-etiolation including those involved in light and chloroplast regulation, as well as the circadian clock. This included two paralogs of the photomorphogenesis master regulator *ELONGATED HYPOCOTYL 5 (HY5)* as well as an ortholog of *EARLY PHYTOCHROME RESPONSIVE 1*. Consistent with other species (Leivar et al. 2008), genes encoding negative regulators of de-etiolation PHYTOCHROME INTERACTING FACTOR7 (PIF7) and PIF3-LIKE5 PIL5 from *G. gynandra* were rapidly down-regulated in light. Orthologs of the chloroplast regulators GNC and GLK2 (Zubo et al. 2018) were also part of these early up-regulated clusters, as were clock components including two paralogs of *REVEILLE (RVE2)*, and one each of *RVE1* and *LATE ELONGATED HYPOCOTYL (LHY)*. In fact, a direct comparison with a de-etiolation time-course from Arabidopsis indicated a striking conservation in the dynamics of genes that were up-regulated in the first 0.5 hours of light exposure.

The principle that C_4_ genes need to become subject to gene regulatory networks that allow co-regulation with other photosynthesis genes is intuitive, however, to our knowledge strong evidence for this phenomenon has been lacking. Our data clearly indicate that in *G. gynandra* transcript abundance of photosynthesis genes increased strongly during de-etiolation. Furthermore, transcripts derived from most C_4_ cycle genes increased over the de-etiolation time course. Re-analysis of publicly available data (Sullivan et al. 2014) showed that, whilst C_4_ genes in Arabidopsis also respond to light, the induction was less strong (Supplemental Figure 3). The most parsimonious explanation for these findings is that in C_3_ plants genes encoding components of the C_4_ pathway show a basal induction in response to light during de-etiolation, and that this ancestral system becomes amplified during the evolution of C_4_ photosynthesis.

### Chromatin dynamics and transcription factor binding during de-etiolation of *G. gynandra*

DNaseI-, MNase- and ATAC-sequencing allow regions of open chromatin to be defined (Klein and Hainer 2020; Tsompana and Buck 2014). In plants, use of such technologies has been initiated to better understand gene expression associated with processes, such as de-etiolation and heat stress (Sullivan et al. 2014), the cold response (Han et al. 2020), and how chromatin accessibility impacts on gene expression across Arabidopsis ecotypes (Alexandre et al. 2018). Defining regions of open chromatin allows motifs available to be bound by transcription factors to be predicted *in silico* (Sullivan et al. 2014). More generally, the identification of accessible regions across a dynamic process offers the chance to identify global chromatin patterns. Previous analysis of leaves from C_3_ and C_4_ plants identified DHS in mature leaves in which photosynthesis had already been established (Burgess et al. 2019). Here, we used de-etiolation to provide insight into how genome-wide patterns of DHS, as well as those specifically associated with photosynthesis genes during assembly of the photosynthetic apparatus. This showed that DHS density around predicted transcription start sites increased during the first two hours of light and is consistent with genes becoming more accessible (Liu et al. 2017). However, global analysis showed that for early responding genes increased DNA accessibility did not always lead to increased gene expression (Supplemental Figure 5). Whilst DHS data alone therefore appear to have limited power to predict gene expression, the ENCODE consortium’s analysis of additional datasets allowed identification of enhancer regions, their interaction with promoters, and relationships between promoters and histone marks and methylation to be elucidated (Thurman et al. 2012). It is therefore probable that deeper insight from DHS will emerge once additional datasets are available.

To start to provide complementary data to those derived from the regions of chromatin that could be bound by transcription factors, we undertook deep sequencing to generate *in vivo* evidence of protein-DNA interactions. Although it has been proposed that this approach can generate base pair resolution information on transcription factor binding sites (Neph et al. 2012), the fact that leaves contain many cell types differing in gene expression likely leads to some DGF representing estimates of binding. Nevertheless, we identified 300,091 DGFs in *G. gynandra* during de-etiolation which compares favourably with 282,030 DGFs in a publicly available dataset for de-etiolation of Arabidopsis (Sullivan et al. 2014) that did not include filtering to remove bias associated with DNaseI activity. Motifs associated with *in vivo* binding in *G. gynandra* supported findings from the DHS analysis above. For example, an increase of binding *in vivo* by bZIP factors was detected at 0.5 and 2 hours after light exposure suggesting a causal link between their induction and the response of early up-regulated genes. Moreover, whilst expression data were used to generate a gene regulatory network that predicted transcription factors belonging to bZIP, homeodomain and TCP families act upstream of C_4_ pathway genes, the DGF data provided *in vivo* evidence for binding of bZIP (G-boxes) and homeodomain motifs in *PPDK* as well as homeodomain and TCP motifs in *BASS2*. Greater sequencing depth as well as analysis of separated mesophyll or bundle sheath cells in the future is likely to provide greater insights into transcription factor binding events associated with the induction and maintenance of C_4_ photosynthesis.

More broadly, genome-wide analysis of DGF showed that between 0 and 2 hours after exposure to light, transcription factor binding decreased in coding regions but increased in promoters and 5′ UTRs. In *Drosophila melanogaster*, enhancers have been reported to be depleted in exons but enriched in promoters, 5′ UTRs, and introns (Arnold et al. 2013). Consistent with this, we found transcription factor binding events predicted to be activators were 1.6 and 2.9 times more likely to be found in promoters and introns respectively, whilst negative regulators were twice as likely to be found in exons. Transcription factor binding sites in coding sequence have been termed duons as they determine both gene expression and the amino acid code. Duons have been proposed to act as repressors of gene expression in humans (Stergachis et al. 2013), and in plants duons placed downstream of the constitutive CaMV35S promoter repress gene expression in mesophyll cells (Brown et al. 2011; Reyna-Llorens et al. 2018). The extent to which duons act as repressors of gene expression remains to be determined, but the data reported here are consistent with this notion.

### Divergence in the C_4_ cistromes of C_3_ Arabidopsis and C_4_ *G. gynandra*

Through analysis of the DHS reported here as well as those from an Arabidopsis dataset associated with de-etiolation (Sullivan et al. 2014) we conclude that the cistromes of C_4_ orthologs in these species are less similar than those of their C_3_ photosynthesis genes. Two phenomena appear to underpin this cistrome divergence. First, C_4_ genes in *G. gynandra* have acquired LREs that are strongly enriched in the C_3_ cistromes of both species. This finding is consistent with C_4_ genes in *G. gynandra* becoming embedded in light regulatory networks. Thus, although C_4_ orthologs in Arabidopsis are under light and chloroplast regulation (Burgess et al. 2016), they appear to have fewer LREs. While LREs in general were enriched in the cistromes associated with photosynthesis genes in both species, GT trihelix motifs represented some of the most preferentially enriched motifs in the *G. gynandra* C_4_ cistrome. It is thus possible that this aspect of light signalling has been disproportionally adopted to regulate the C_4_ pathway. Divergence between the cistromes of C_4_ genes from Arabidopsis and *G. gynandra* was also caused by more TCP, homeodomain and YABBY binding sites in the C_4_ species. While these motifs seem to link many C_4_ genes to early acting components of the de-etiolation network, their prevalence in the *G. gynandra* C_4_ cistrome implies a particular role associated with the C_4_ pathway. One possibility is that these motifs are involved in the cell gene expression patterning required to establish the carbon pump between mesophyll and bundle sheath cells. However, at the moment it is unclear if these changes in *cis* allow the C_4_ genes to be rewired such that they become connected to regulatory networks that operate in mesophyll or bundle sheath cells of the C_3_ leaf. As only one example associated with patterning genes to the Arabidopsis bundle sheath has been documented to date (Dickinson et al. 2020), it is possible that members of the TCP, homeodomain and YABBY families which are over-represented in C_4_ genes of *G. gynandra* are also involved in re, stricting gene expression to these cell types.

### C_4_ genes in *G. gynandra* are connected to the top of de-etiolation gene networks

Coupling of RNA- and DNaseI-SEQ provided insight into how the C_4_ pathway is regulated during de-etiolation but also how it has likely evolved. As *G. gynandra* is in a family sister to the Brassicaceae, the function of transcription factor orthologs can be inferred from Arabidopsis. Consistent with this, the response of gene expression during de-etiolation in *G. gynandra* was similar to Arabidopsis with many of the master regulators of photomorphogenesis and chloroplast development as well as components of the circadian clock being among the first transcripts induced after exposure to light (Figure 2E-F and Supplemental Table 2). This included two paralogs of HY5 that showed similar expression dynamics to each other and to their putative ortholog in Arabidopsis. HY5 acts at the top of the transcriptional hierarchy and is considered a “master regulator” directly binding thousands of targets (Lee et al. 2007). It also integrates a number of inputs including light through its interaction with COP1 and subsequent degradation (Osterlund, Wei, and Deng 2000), as well as hormone pathways including inhibition of auxin signalling to suppress hypocotyl elongation (Cluis, Mouchel, and Hardtke 2004), and repression of ethylene signalling through the activation of the repressor ERF11 (Li et al. 2011).

Regulatory networks are complex but have general structures that include hierarchy and modularity. The hierarchy associated with activation of photosynthesis gene expression in *G. gynandra* indicates components that likely act by 0.5 hours to modify downstream dependent components. As cotyledons of *G. gynandra* operate C_4_ photosynthesis, it is logical that induction of C_4_ and C_3_ photosynthesis genes are both linked to the top of the regulatory networks associated with de-etiolation. We detected evidence of this integration from multiple orthogonal analyses including the predicted regulation of C_4_ pathway components by transcription factors that themselves are highly connected in the transcription factor regulatory network. One example was the predicted regulation of the Phospho*enol*pyruvate/Phosphate Translocator (PPT) by TCP8 (Figure 3B). TCP8 was also one of the most highly connected transcription factors in the network with multiple in- and out-edges suggesting a central role integrating early regulation during the first few hours of de-etiolation. By integrating analysis of transcript abundance with *in-silico* analysis of cistromes we identified motifs enriched in DHSs of many photosynthesis genes up-regulated at 0.5 hours. This showed a strong association with bZIP (G-box), bHLH (E-box), MYB-related (I-box) and GT trihelix motifs linked to light signalling (Chattopadhyay et al. 1998; Gangappa and Botto 2016; Gilmartin and Chua 1990), CCA1-related clock motifs as well as homeodomain and YABBY motifs. A strong overlap was detected between these motifs associated with photosynthesis genes and those most strongly enriched in the C_4_ gene cistrome. Furthermore, *in vivo* evidence of binding to G-boxes was detected in *PPDK*. We conclude that a range of transcription factors at the top of the de-etiolation regulatory network rapidly induce the gene expression needed for both C_3_ and C_4_ photosynthesis. Notably, our data do not indicate an obvious master regulator of C_4_ photosynthesis (Westhoff and Gowik 2010) but rather that the genes have become integrated with existing regulatory mechanisms including canonical light regulation networks.

Some transcription factors implicated in early de-etiolation and C_4_ regulation function outside of light signalling. For example, YABBY motifs were enriched in genes up-regulated genes at 0.5 hours, but also the C_4_ cistrome, and the YABBY AFO/FIL transcription factor was predicted to act as a regulator of *PPa6*. Although in mature leaves YABBY transcription factors are implicated in auxin patterning (Sarojam et al. 2010), in the developing leaf they are important in specifying abaxial cell fate (Siegfried et al. 1999). AFO/FIL also positively regulates glucosinolate biosynthesis (Douglas et al. 2017) for which gene expression in the bundle sheath is conditioned (Aubry et al. 2014). It therefore seems possible that YABBY factors play a role in patterning genes to the bundle sheath of C_4_ *G. gynandra*.

Homeodomain factors were identified as likely early regulators of de-etiolation as well as of C_4_ genes (Figure 4D and H). For example, in the cistrome of genes up-regulated at 0.5 hours motifs from both the HD-ZIPI and HD-ZIPII families were over-represented. Furthermore, the gene regulatory network determined form expression data modelling predicted that *PEPC* is regulated by the HD-ZIPII HB-2. HB-2 has previously been implicated in the shade response and cotyledon expansion (Steindler et al. 1999). Another HD-ZIPII factor, HAT3, was also highly connected to the de-etiolation transcription factor network of *G. gynandra*. Members of the HD-ZIP families are involved in a wide range functions from vascular development (e.g., HD-ZIPIII ATHB8 and ATH15 (Baima et al. 2001; Ohashi-Ito and Fukuda 2003) to anthocyanin production e.g., HD-ZIPIV ANL2 (Kubo et al. 1999). It is possible that they have aquired additional roles that link more closely to photosynthesis in the C_4_ leaf. Of the enriched homeodomain motifs present in genes induced at 0.5 hours but also in the C_4_ cistromes, the most common were those bound by HD-ZIPI factors. Two HD-ZIPI factors predicted in the gene regulatory network, HB-7 (Valdés et al. 2012) and HB20 (Barrero et al. 2010) are implicated in ABA signalling. However, given the similarity in their DNA recognition sequence, it is not clear which HD-ZIPs regulate C_4_ pathway genes.

TCP motifs were most strongly enriched in genes upregulated after two hours of light and were also found to peak in the DGF data at the same time point. As TCP motifs were also among the fifty most enriched motifs in the C_4_ cistrome, components of the C_4_ cycle may be under their direct control. The closely related Class II TCPs, TCP7 and TCP8, were central to the gene regulatory network with many in- and out-edges and TCP8 was predicted to regulate *PPT*. To date TCPs have been linked to a number of functions and through promiscuous binding interact with a wide range of other transcription factor families (Trigg et al. 2017). TCP7 and TCP8 are expressed strongly in emerging leaves where they impact on leaf shape and number through cell proliferation (Aguilar-Martínez and Sinha 2013). As the developing cotyledon undergoes cell proliferation and expansion, it is possible that TCP7 and TCP8 regulate these processes during early cotyledon development, and that components of the C_4_ cycle have been rewired to link to this network.

Perhaps the clearest outputs from our analysis of de-etiolation and C_4_ gene regulation in *G. gynandra* relate to light signalling. Light receptors such as phytochromes, cryptochromes, phototropins and UV-B receptors function co-operatively to allow the integration of light cues (Kong and Okajima 2016). Their signalling terminate at a number of LREs to guide the transcriptional machinery to target genes. LREs including G-, E-, I- and GT-boxes are bound by members of the bZIP (Foster, Izawa, and Chua 1994), bHLH (Yadav et al. 2005) and BZR (Yin et al. 2005), MYB-related (Rose, Meier, and Wienand 1999) and trihelix GT factors (Green, Kay, and Chua 1987) respectively. *In vivo* binding to GT-and G-boxes to *PPDK* provides direct evidence that such sites are bound during assembly of the photosynthetic apparatus. The acquisition of LREs by C_4_ genes represents a simple conceptual route to allow them to be co-regulated with photosynthesis gene expression. Although it has been proposed that C_4_ orthologs in Arabidopsis are controlled by light and chloroplast signalling (Burgess et al. 2016) to our knowledge, there is little direct evidence relating to the extent to which this ancestral regulation is modified during the evolution of C_4_ photosynthesis. The data presented here argue for some C_4_ genes being strongly responsive to light because they contain multiple copies of the same motif whereas others have acquired multiple different LREs. This is consistent with the fact that the pea *RbcS* promoter contains multiple LREs (Gilmartin and Chua 1990), and suggests that for genes to become highly light responsive they require more than one type of LRE (Chattopadhyay et al. 1998). Taken together, our findings suggest that C_4_ genes become linked to other photosynthesis genes via changes in *cis* such that they are responsive to ancestral gene regulatory networks downstream of light. This integration of C_4_ genes into ancestral gene regulatory networks should allow rapid, robust, and spatially precise induction during de-etiolation pf the C_4_ leaf.

## MATERIALS AND METHODS

### Plant growth, chlorophyll quantitation and microscopy

G*ynandropsis gynandra* seeds were sown directly from intact pods and germinated on moist filter papers in the dark at 32 °C for 24 hours. Germinated seeds were then transferred to half strength Murashige and Skoog (MS) medium with 0.8 % (w/v) agar (pH 5.8) and grown for three days in a growth chamber at 26 °C. De-etiolation was induced by exposure to white light with a photon flux density (PFD) of 350 μmol m^-2^ s ^-1^ and photoperiod of 16 hours. Whole seedlings were harvested at 0.5, 2, 4 and 24 hours after illumination (starting at 8:00 with light cycle 6:00 to 22:00). Tissue was flash frozen in liquid nitrogen and stored at −80 °C prior to processing.

For analysis of chlorophyll content, de-etiolating *G. gynandra* seedlings were flash frozen at 0, 0.5, 2, 4 or 24 hours post light exposure. 100 mg of tissue was suspended in 1 ml 80 % (v/v) acetone at 4 °C for 10 minutes prior to centrifugation at 15,700 g for 5 minutes and removal of the supernatant. The pellet was resuspended in 1 ml 80 % (v/v) acetone at 4 °C for 10 minutes, precipitated at 15,700 g for 5 minutes. Supernatants were pooled, and absorbance measured in a spectrophotometer at 663.8 nm and 646.6 nm. Total chlorophyll content determined as described previously (Porra, Thompson, and Kriedemann 1989). For electron microscopy, *G. gynandra* cotyledons (∼2 mm^2^) were excised with a razor blade and fixed immediately in 2 % (v/v) glutaraldehyde and 2 % (w/v) formaldehyde in 0.05-0.1M sodium cacodylate (NaCac) buffer (pH 7.4) containing 2 mM calcium chloride. Samples were vacuum infiltrated overnight, washed five times in deionized water, and post-fixed in 1 % (v/v) aqueous osmium tetroxide, 1.5 % (w/v) potassium ferricyanide in 0.05□M NaCac buffer for 3 days at 4 °C. After osmication, samples were washed five times in deionized water and post-fixed in 0.1 % (w/v) thiocarbohydrazide in 0.05□M NaCac buffer for 20 minutes at room temperature in the dark. Samples were then washed five times in deionized water and osmicated for a second time for 1 hour in 2 % (v/v) aqueous osmium tetroxide in 0.05□M NaCac buffer at room temperature. Samples were washed five times in deionized water and subsequently stained in 2 % (w/v) uranyl acetate in 0.05 M maleate buffer (pH 5.5) for 3 days at 4 °C and washed five times afterwards in deionized water. Next, samples were dehydrated in an ethanol series, transferred to acetone, and then to acetonitrile. Samples were embedded in Quetol 651 resin mix (TAAB Laboratories Equipment Ltd). For transmission electron microscopy (TEM), ultra-thin sections were cut with a diamond knife, collected on copper grids and examined in a FEI Tecnai G2 transmission electron microscope (200 keV, 20 µm objective aperture). Images were obtained with AMT CCD camera. For scanning electron microscopy (SEM), ultrathin-sections were placed on plastic coverslips which were mounted on aluminium SEM stubs, sputter-coated with a thin layer of iridium and imaged in a FEI Verios 460 scanning electron microscope. For light microscopy, thin sections were stained with methylene blue and imaged by an Olympus BX41 light microscope with a mounted Micropublisher 3.3 RTV camera (Q Imaging).

### RNA and DNaseI sequencing

Before processing, frozen samples were divided into two, the first being used for RNA-SEQ analysis and the second for DNaseI-SEQ. Samples were ground in a mortar and pestle and RNA extraction carried out with the RNeasy Plant Mini Kit (74904; QIAGEN) according to the manufacturer’s instructions. RNA quality and integrity were assessed on a Bioanalyzer High Sensitivity DNA Chip (Agilent Technologies). Library preparation was performed with 500 ng of high integrity total RNA (RNA integrity number > 8) using the QuantSeq 3’ mRNA-SEQ Library Preparation Kit FWD for Illumina (Lexogen) following the manufacturer’s instructions. Library quantity and quality were checked using Qubit (Life Technologies) and a Bioanalyzer High Sensitivity DNA Chip (Agilent Technologies). Libraries were sequenced on NextSeq 500 (Illumina, Chesterford, UK) using single-end sequencing and a Mid Output 150 cycle run.

To extract nuclei, tissue was ground in liquid nitrogen and incubated for five minutes in 15mM PIPES pH 6.5, 0.3 M sucrose, 1 % (v/v) Triton X-100, 20mM NaCl, 80 mM KCl, 0.1 mM EDTA, 0.25 mM spermidine, 0.25 g Polyvinylpyrrolidone (SIGMA), EDTA-free proteinase inhibitors (ROCHE), filtered through two layers of Miracloth (Millipore) and pelleted by centrifugation at 4 °C for 15 min at 3600 g. To isolate deproteinated DNA, 100 mg of tissue from seedlings exposed to 24 hours light were harvested two hours into the light cycle, four days after germination. DNA was extracted using a QIAGEN DNeasy Plant Mini Kit (QIAGEN, UK) according to the manufacturer’s instructions. 2×10^8^ nuclei were re-suspended at 4 °C in digestion buffer (15 mM Tris-HCl, 90 mM NaCl, 60 mM KCl, 6 mM CaCl_2_, 0.5 mM spermidine, 1 mM EDTA and 0.5 mM EGTA, pH 8.0). DNAse-I (Fermentas) at 2.5 U was added to each tube and incubated at 37 °C for three minutes. Digestion was arrested by adding a 1:1 volume of stop buffer (50 mM Tris-HCl, 100 mM NaCl, 0.1 % (w/v) SDS, 100 mM EDTA, pH 8.0, 1 mM Spermidine, 0.3 mM Spermine, RNaseA40 µg/ml) and incubated at 55 °C for 15 minutes. 50 U of Proteinase K were then added and samples incubated at 55 °C for 1 h. DNA was isolated by mixing with 1 ml 25:24:1 Phenol:Chloroform:Isoamyl Alcohol (Ambion) and spun for 5 minutes at 15,700 g followed by ethanol precipitation of the aqueous phase. Samples were size-selected (50-400 bp) using agarose gel electrophoresis and quantified fluorometrically using a Qubit 3.0 Fluorometer (Life technologies), and a total of 10 ng of digested DNA (200 pg l^-1^) used for library construction. Sequencing ready libraries were prepared using a TruSeq Nano DNA library kit according to the manufacturer’s instructions. Quality of libraries was determined using a Bioanalyzer High Sensitivity DNA Chip (Agilent Technologies) and quantified by Qubit (Life Technologies) and qPCR using an NGS Library Quantification Kit (KAPA Biosystems) prior to normalisation, and then pooled, diluted and denatured for paired-end sequencing using High Output 150 cycle run (2x 75 bp reads). Sequencing was performed using NextSeq 500 (Illumina, Chesterford UK) with 2x 75 cycles of sequencing.

### RNA-SEQ data processing and quantification

Commands used are available on GitHub (“command_line_steps”) but an outline of steps was as follows. Raw single ended reads were trimmed using trimmomatic (Bolger, Lohse, and Usadel 2014) (version 0.36). Trimmed reads were then quantified using salmon (Patro et al. 2017) (version 0.4.234) after building an index file for a modified *G. gynandra* transcriptome. The transcriptome was modified to create a pseudo-3’ UTR sequence of 339 bp (the mean length of identified 3’UTRs) for *G. gynandra* gene models that lacked a 3’ UTR sequence which was essentially an extension beyond the stop codon of the open reading frame. Inclusion of this psuedo 3’ UTR improved mapping rates. Each sample was then quantified using the salmon “quant” tool. All *.sf files had the “NumReads” columns merged into a single file (All_read_counts.txt) to allow analysis with both DEseq2 (Anders and Huber 2010) and edgeR (McCarthy, Chen, and Smyth 2012). The edgeR pipeline was run as the edgeR.R R script (on GitHub) on the All_read_counts.txt file to identify the significantly differentially expressed genes by comparing each time-point to the previous one. A low expression filter step was also used. We then similarly analysed the data with the DEseq2 package using the DEseq2.R R script (on GitHub) on the same All_read_counts.txt file. This also included the PCA analysis shown in Figure 2A. The intersection from both methods was used to identify a robust set of differentially regulated genes. For most further analysis of the RNA-SEQ data, mean TPM values for each time-point (from three biological replicates) was first quantile normalised and then each value divided by the sample mean such that a value was of 1 represented the average for that sample. This processing facilitated comparisons between experiments across species in identifying changes to transcript abundance between orthologs.

### GO enrichment analysis, identification of C_3_ and C_4_ gene lists and heatmap plotting

The agrigo-v2 web tool was used for GO analysis following the tools instructions for a custom background. The background was made by mapping all *G. gynandra* genes to their closest Arabidopsis blastp hit and inheriting all the GO terms associated with that gene from the TAIR gene annotation file (Athaliana_167_TAIR10.annotation_info.txt from Phytozome). Differentially expressed genes from each time-point were analysed and GO terms with significance < 10^−5^ in at least one differentially expressed gene set were kept (Supplemental Table 1, Supplemental Figure 2). Representative GO terms were selected for plotting in a stacked barplot using the R script (Fig2B.R) and data file Fig2B_GO_term_data.txt (on GitHub).

In order to map orthologs between Arabidopsis and *G. gynandra*, OrthoFinder (Emms and Kelly 2019) was used. This allows more complex relationships than a 1:1 to be identified and placed into orthogroups. C_3_ photosynthesis genes were first identified from Arabidopsis through the “photosynthesis” (GO:0015979) keyword search on the TAIR browse tool (https://www.arabidopsis.org/servlets/TairObject?type=keyword&id=6756) and resulted in ninety-two genes for Arabidopsis. Their orthologs were found in *G. gynandra* using the orthogroups generated between the two species and resulted in ninety-three C_3_ photosynthesis genes. C_4_ genes used in this study are considered the “core” pathway genes and are a manually curated set largely based on previous analysis (Burgess et al. 2016). Initially, multiple paralogs were included but non-induced transcripts were then filtered out. Orthologs between the two species were again identified from the orthogroups from OrthoFinder. The *G. gynandra* C_3_ and C_4_ gene normalised expression values were further processed with each value being divided by the row mean and log10(x/row mean) plotted as a heatmap using the R script Fig2C.R on data file Fig2C_heatmap_data.txt (on GitHub). The heatmap for Arabidopsis C_3_ and C_4_ gene expression was made in the same way as for the *G. gynandra* data using the gene lists as previously described.

To generate the heatmap of all highly variable transcription factors, all putative transcription factors were extracted from the DEGs. All potential transcription factors in *G. gynandra* were obtained by blastp searching with all protein sequences from the Arabidopsis Plant Transcription Factor Database (http://plntfdb.bio.uni-potsdam.de/v3.0/downloads.php?sp_id=ATH) with command available on GitHub. All predicted transcription factors found in the DEGs had their normalised TPM values converted to z-scores (by row) and plotted using the heatmap.2 R package using the Fig2D.R script with the Fig2D_TF_Expression_Data.txt file. Clusters were manually assigned from the hierarchical clustering and word clouds made of transcription factor family names for each cluster. Data from clusters IV and V was plotted as boxplots alongside orthologs normalised expression values from Arabidposis de-etiolation data using the Fig2E-F.R script on Fig2E_data.txt and Fig2F_data.txt.

### Time series Gene Network Construction

To build the transcription factor network, all predicted *G. gynandra* transcription factors had all three replicate expression values from 0, 0.5, 2 and 4 hours selected and then filtered to remove those with low expression and low variance. The resulting 290 transcription factors were used as input for both the dynGENIE3, using default settings, and Sliding Window Inference for Network Generation (SWING), using kmin and kmax of 1 and a window (w) of 3, tools following user instructions. The SWING settings allow the model to consider a single step cause and effect delay between the windows of 0, 0.5 and 2 hours and 0.5, 2 and 4 hours. Following construction of the two networks, the edge weights (predicted strength of interaction) were added together (Combined_SWING_dynG3.txt on GitHub). The resultant network was filtered based on the edge weights such that all nodes were still represented with at least one edge. This network was then loaded into Cytoscape and a final “core” network was selected for visualisation by keeping only those nodes with the most edges (in and out) as well as the highest betweenness centrality. To predict regulators of the C_4_ pathway genes, these were added to the pipeline and analysed as described above. The final network for visualisation was made up of the highest confidence regulators (only out-edges) of the C_4_ pathway genes (only in-edges).

### DNaseI-SEQ data processing

Three biological replicates for each time-point were sequenced in multiple runs with one sample being chosen, based on initial QC scores, for deeper sequencing to provide the necessary depth for calling both DNaseI Hypersensitive Site (DHS) and Digital Genomic Footprints (DGF). For each sample, raw reads from multiple sequencing runs were combined and trimmed for low quality reads using trimmomatic. These files were analysed with fastqc (http://www.bioinformatics.babraham.ac.uk/projects/fastqc/) to ensure samples passed QC parameters (see Ggynandra_DNaseSEQ_multiqc_report.html on GitHub) and then mapped to the *G. gynandra* genome using bowtie2 (version 2.3.4.1) with the “--local” pre-set option. Following mapping, a bash script (DNaseSEQ_tagAlign.sh on GitHub) was run on each bam file which in summary: filters low quality (MAPQ < 30) mapped reads and plots MAPQ distribution, removes duplicates, measures library complexity, fragment sizes, GC bias and finally makes tagAlign files. The three tagAlign files from each time-point were then merged before running another bash script (DHS_DGF_identification.sh on GitHub) for each time-point which in summary: uses “macs2 callpeak” to identify “narrowPeaks”, finds the distance for each DHS to its closest transcriptional start site (TSS), calculates SPOT scores (see https://www.encodeproject.org/data-standards/dnase-SEQ/), plots DHS profiles using the deeptools bamCoverage, computeMatrix and plotProfile tools and finally calls DGF using the wellington_footprints.py program (Piper et al. 2013). The footprints identified with a log(*p*-value) cut-off of < −10 were used for further analysis.

In order to carry out comparative analysis between *G. gynandra* and Arabidopsis, an analogous de-etiolation time-course (Sullivan et al. 2014) was reprocessed in the same way as the *G. gynandra* data. Arabidopsis DNaseI-SEQ data was mapped to the TAIR9 genome and RNA-SEQ was mapped using Salmon to the Araport11 transcriptome, followed by the use of tximport to collapse expression values for all isomers into a single value, a step not required for *G. gynandra* as it lacks isomer information. To allow inter-species comparisons, as with *G. gynandra* the Arabidopsis RNA-SEQ data were quantile normalised and then each value divided by the samples mean expression value.

### Analysis of DHS changes across the time-course

To plot the position of DHS relative to TSS regions, the center of each DHS was found and the distance measured to the nearest TSS site using “bedtools closest”. These distances were then plotted as a density plot for each time point using Fig4B.R script with the Fig4B_DHS_TSS_data.txt.

To quantify the overlap in DHS between samples, DHS (from the “narrowPeak” file) were sorted by their “-log10qvalue” column and only the top ranked DHS regions used until a total of 55,122,108 bp was reached which corresponded to the total length of DHS regions in the 4 hours sample which had the least. This allowed us to compare overlap between equal sized regions. These DHS regions for each time-point were then intersected in a pairwise fashion using bedtools intersect. The total length of intersecting regions was divided by 55,122,108 bp to obtain the proportion of overlap for each pairwise comparison generating the values in Figure 4C.

To compare the differential DHS (dDHS) scores for gene sets of interest, we defined promoter regions around each gene of interest as 1000 bp upstream from predicted transcription start site. We then identified DHS regions that intersected with each gene of interest (i.e., all DHS regions overlapping with a gene body or promoter) and merged these DHS regions (equivalent to an outer join). The dDHS tool (He et al. 2012) as part of the pyDNase package (Piper et al. 2015) was used to quantify changes in accessibility between consecutive time-points for a given region, where SAMPLEA precedes SAMPLEB in the time-course. Finally, these dDHS values were plotted in violin plots.

### Motif Analysis of DHS regions

To identify enriched motifs in the cistromes of the DEGs from the 0.5, 2 and 4 hour time points, we first scanned for the presence of motifs from the DAPseq (O’Malley et al. 2016) and PBM (Franco-Zorrilla et al. 2014) databases using the meme suite FIMO tool (Grant, Bailey, and Noble 2011) in all DHS regions that intersected with low expression filtered gene loci (gene body or 1.5kbp promoter region). This formed a background from which permutation testing was carried out using the regioneR R package (Gel et al. 2016). In summary, for each set of DEGs, the intersecting “n” DHS had the motif frequencies across all regions compared with >1000 bootstraps of “n” randomly sampled regions to generate a probability of the observed frequencies being from the null hypothesis of no enrichment. These p-values were then Bonferroni corrected for the number of motifs tested (n=937) with any motifs passing the new ∼99.995% confidence interval defined as enriched (or depleted). We selected motifs that were only enriched with either the up- or down-regulated DHS of each time-point and further collapsed these down by motif groups given the high similarity of many related motifs (i.e., they match the same regions).

In order to allow a comparison between the cistromes of the C_3_ and C_4_ genes from both *G. gynandra* and Arabidopsis, we first filtered out those genes that did not show induction across the time-course (red clusters from photosynthesis heatmaps). We then found the fold change of observed motifs frequencies in their DHS regions versus a random set (from 1500 random genes) for each species. These values were then ranked with 1 representing the most strongly “enriched” motif and the rankings compared. To analyse and compare motifs between species we ranked motifs by their normalised frequencies against the background for each gene sets DHS (C_4_ pathway and C_3_ photosynthesis from both *G. gynandra* and *A. thailiana*). These sets were filtered to remove genes that were not induced during the time-course. The top 50 motifs from each set were then plotted against their rank in other sets. As motifs that were highly ranked in *G. gynandra* C_4_ genes were of particular interest, these were plotted as a heatmap using the Fig4H.R R script on the Fig4E-H_data file (on GitHub).

### DNaseI bias correction

To reduce the proportion of false positive DGF calls caused by DNAseI cutting bias, DNaseI-SEQ was performed on de-proteinated gDNA and mapped to the *G. gynandra* genome. The hexamer cutting frequencies at the DNaseI cutting sites were used to generate a background signal profile that was incorporated into a mixture model to calculate the log-likehood ratio (FLR) for each footprint using the R package MixtureModel (Yardimci et al. 2014). DGF with low confidence (FLR<0) were filtered out resulting in a reduction of 11 to 37 % of DGF per timepoint. Same pipeline was used in previous analysis (Burgess et al. 2019). The pipeline is illustrated in Supplemental Figure 4.

### DGF genomic feature distributions DGF motif frequencies and DGF-target correlation

To identify the distribution of DGF across genomic features we used bedtools intersect to find the frequency of intersection between the DGF with features in the genome annotation gff3 file promoter (2000 bp upstream of TSS), 5’ UTR, CDS, intron, 3’ UTR and intergenic). These frequencies were divided by the total length of each feature across the genome to determine a density of DGF per feature and these values were plotted as a pie-chart for each time point. In order to link DGF to possible transcription factors, we scanned all DGF using the meme suite fimo tool for both DAPseq and PMB motifs as described for DHS analyses. To visualise how motif frequencies changed during de-etiolation, the frequency of each motif at each time-point was first normalised by the total number of motifs at that time point and then each value was mean centred across the time-course for plotting as a heatmap using the Fig5D.R R script on the Fig5D_heatmap_data.txt file (on GitHub). This hierarchical clustering was then manually grouped and word clouds generated for the motif transcription factor families using an online tool (https://www.wordclouds.com/) for each motif cluster.

Once individual DGF were annotated with potential motifs, we correlated the changes in frequency of each motif with the mean expression of all potential targets. The changes in frequency of each motif were used as a proxy for transcription factor abundance and/or activity and potential targets identified as those genes lying closest to a DGF with a specific motif. A strong positive correlation was used to suggest positive regulation while a strong negative correlation suggests inhibitory regulation. Once potential strong positive and negative regulators were identified, we calculated the percentage of DGF in the different gene features and carried out a Chi goodness of fit test on these proportions based on the null hypothesis that there was no difference in proportions of gene features.

### Generation of C_4_ gene schematics

To create the tracks for schematics, the fimo hits were filtered based on those that showed enrichment in the 0.5 up-regulated gene cistromes (“de-etiolation motifs”) and those in the top 50 most enriched motifs from the *G. gynandra* C_4_ cistrome (“C_4_ motifs”). These bed files were then loaded onto Integrated Genome Viewer (IGV) alongside the bed file for the DGF. Resultant images were exported as SVG files.

## ACCESSION NUMBERS

Raw sequencing data files are deposited in The National Center for Biotechnology Information (PRJNA640984). For full methods, commands, and scripts, see GitHub (https://github.com/hibberd-lab/Singh-Stevenson-Gynandra).

## ACKNOWLEDGMENTS

We thank the Cambridge Advanced Imaging Centre (CAIC) at the University of Cambridge for their support and assistance with electron microscopy, and Anoop Tripathi for acquiring images of greening cotyledons. The work was supported by European Research Council Grant 694733 Revolution, BBSRC Grant BBP0031171 to JMH. GR was supported by a Gates Cambridge Trust PhD Student Fellowship and TBS by Swiss National Science Foundation.

## AUTHOR CONTRIBUTIONS

PS, SRS and JMH designed the study. PS carried out the experimental work. SRS and PS analysed the data. IRL performed de-proteinated DNaseI data analysis. GR assisted in DNaseI assays and library preparations. TBS and PS carried out electron microscopy. PS, SRS and JMH wrote the article and prepared the figures.

## SUPPLEMENTAL FIGURE LEGENDS

**Supplemental Figure 1:** Representative light and scanning electron microscope (SEM) images at 0 and 24 hours after exposure to light. Scale bars represent 100 µm for light microscope images, and 500 nm for SEM.

**Supplemental Figure 2:** GO term enrichment analysis for differentially expressed genes compared with each previous time point. Significantly enriched GO terms were identified using AgriGov2 using a custom *G. gynandra* background built by mapping *G. gynandra* proteins to their closest match in Arabidopsis and inheriting their terms from the TAIR10 annotations. Values plotted are −log10(FDR) and values derived from the up-regulated gene sets shown in red while those form the down-regulated are shown in blue. Many light and photosynthesis-related terms were enriched in the 0.5 hours up-regulated genes. Many primary and secondary metabolism terms were enriched in the 24 hours up-regulated genes.

**Supplemental Figure 3:** Expression patterns of photosynthesis genes (grey sidebar) and C_4_ genes (black sidebar) of *A. thaliana* during de-etiolation. Heatmap illustrating gene expression with each gene being represented by a row, and data centred around the row mean. Dendrograms (red, yellow and green) highlight distinct expression clusters representing strong, moderate or no induction respectively.

**Supplemental Figure 4:** Pipeline for DNaseI-SEQ data processing. On the top left-hand side, pooled reads went through quality control before DHS identification MACS2 peakcalling. The DHSs, representing accessible chromatin regions, were then searched for DGF using the pyDNase package. These DGF are prone to distorting effects due to DNaseI bias in gDNA digestion. On the top right, the pipeline for identifying this bias is shown including the DNaseI digestion of deproteinised (“naked”) gDNA. This generates 6-mer frequencies at each cut site which is used as input for the FootPrintMixture.R tool which scores the Footprint Likelihood Ratio (FLR) of each DGF (likelihood of being a true positive). DGF with FLR < 0 were removed leaving a final set of DGF which were used for analysis. Heatmap of cut patterns are shown centred around each DGF.

**Supplemental Figure 5:** Violin plots showing the distribution of dDHS scores for DHS overlapping with differentially expressed genes at 0.5 hours with both up-regulated (A) and down-regulated shown (B). Mean values are shown as line. Positive and negative dDHS scores represent an increase and decrease in DHS accessibility respectively. No clear association was observed with upregulated C_4_ genes and positive dDHS values or down-regulated genes with negative dDHS values. C. Violin plots depicting changes in DHS accessibility (dDHS) associated with photosynthesis genes, C_4_ photosynthesis genes. Changes are relative to the previous timepoint, n values are for the number of DHS regions quantified.

**Supplemental Figure 6:** DGF in C_4_ cycle genes during de-etiolation time course in *G. gynandra*.

