## Supplementary material for "Induction of C_4_ genes evolved through changes in *cis* allowing integration into ancestral C_3_ gene regulatory networks": S Fig 1

### Supplemental Figure 1

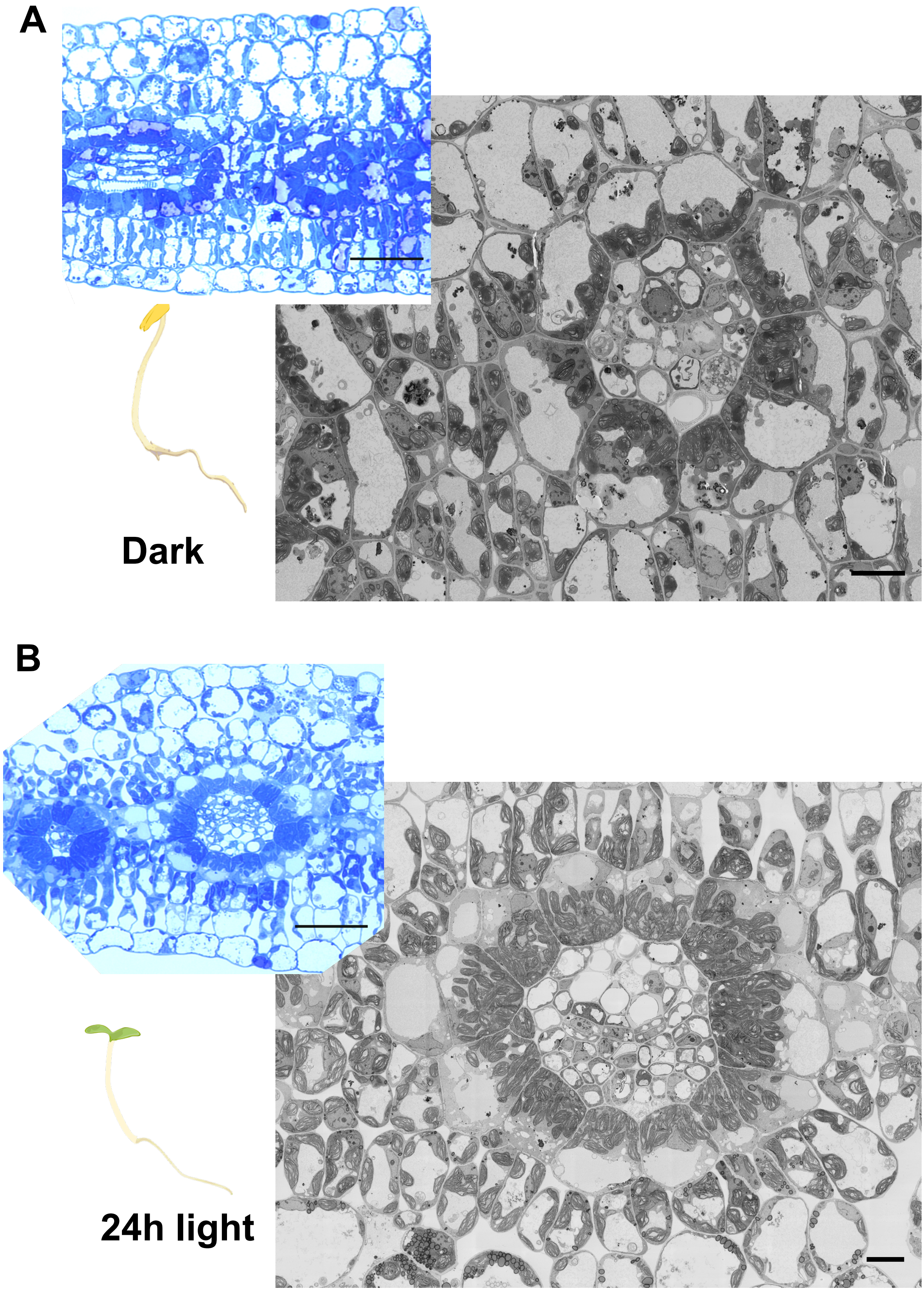


**Supplemental Figure 1**: Representative light and scanning electron microscope (SEM) images of 0 hours (A) and 24 hours (B) de-etiolating *G. gynandra* seedlings at 0 and 24 hours after exposure to light. Scale bars represent 100 µm for light microscope images, and 500 nm for SEM.

### Supplemental Figure 2

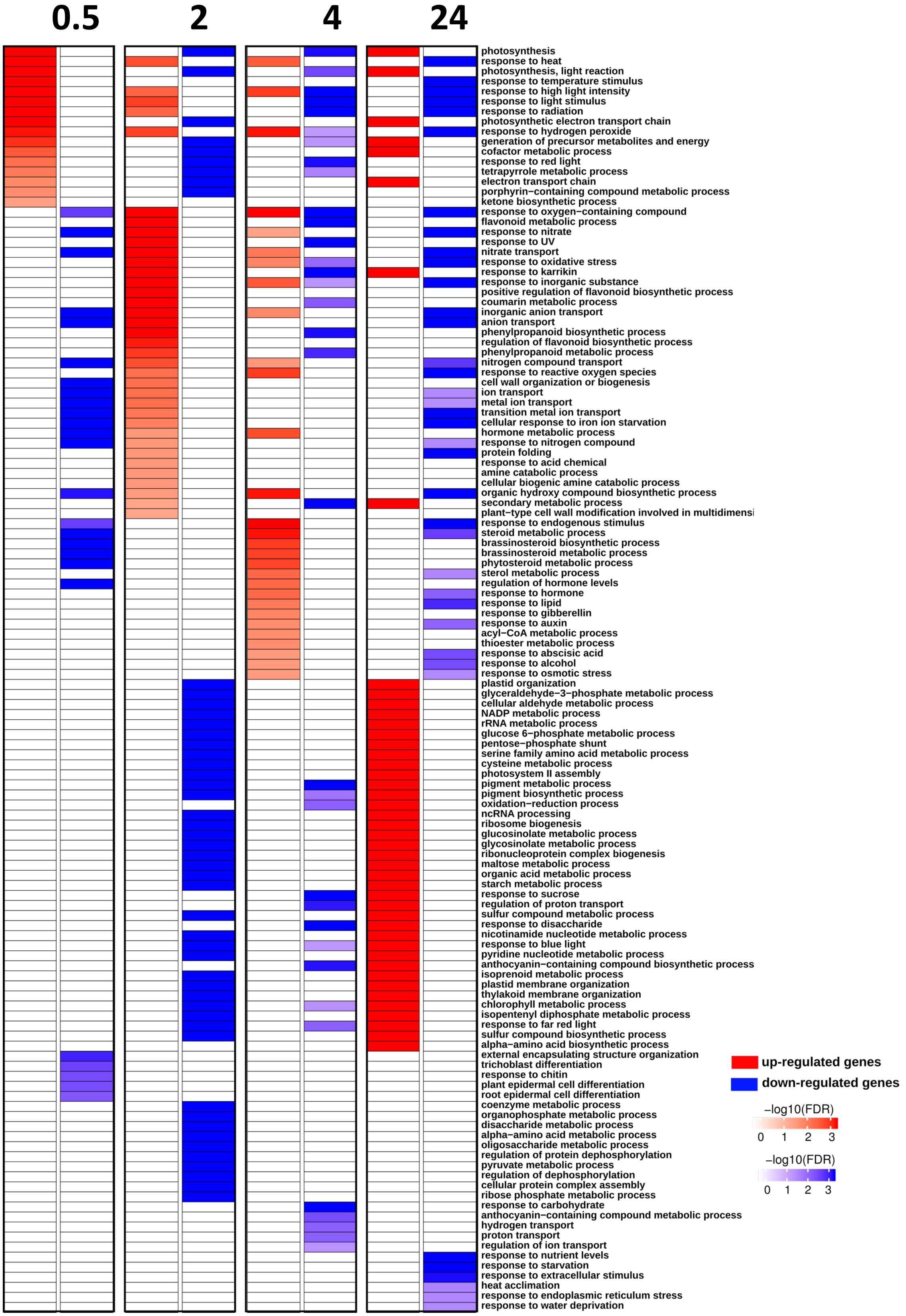


**Supplemental Figure 2**: GO term enrichment analysis for differentially expressed genes as compared to the previous time point. Significantly enriched GO terms were identified using AgriGov2 using a custom *G. gynandra* background built by mapping *G. gynandra* proteins to their closest match in Arabidopsis and inheriting their terms from the TAIR10 annotations. Values plotted are -log10(FDR) and values derived from the up-regulated gene sets are shown in red while those form the down-regulated are shown in blue. Many light and photosynthesis-related terms are enriched in the 0.5 hours up-regulated genes. Many primary and secondary metabolism terms are enriched in the 24 hours up-regulated genes suggesting that photosynthates are being produced by the end of the time course.

### Supplemental Figure 3

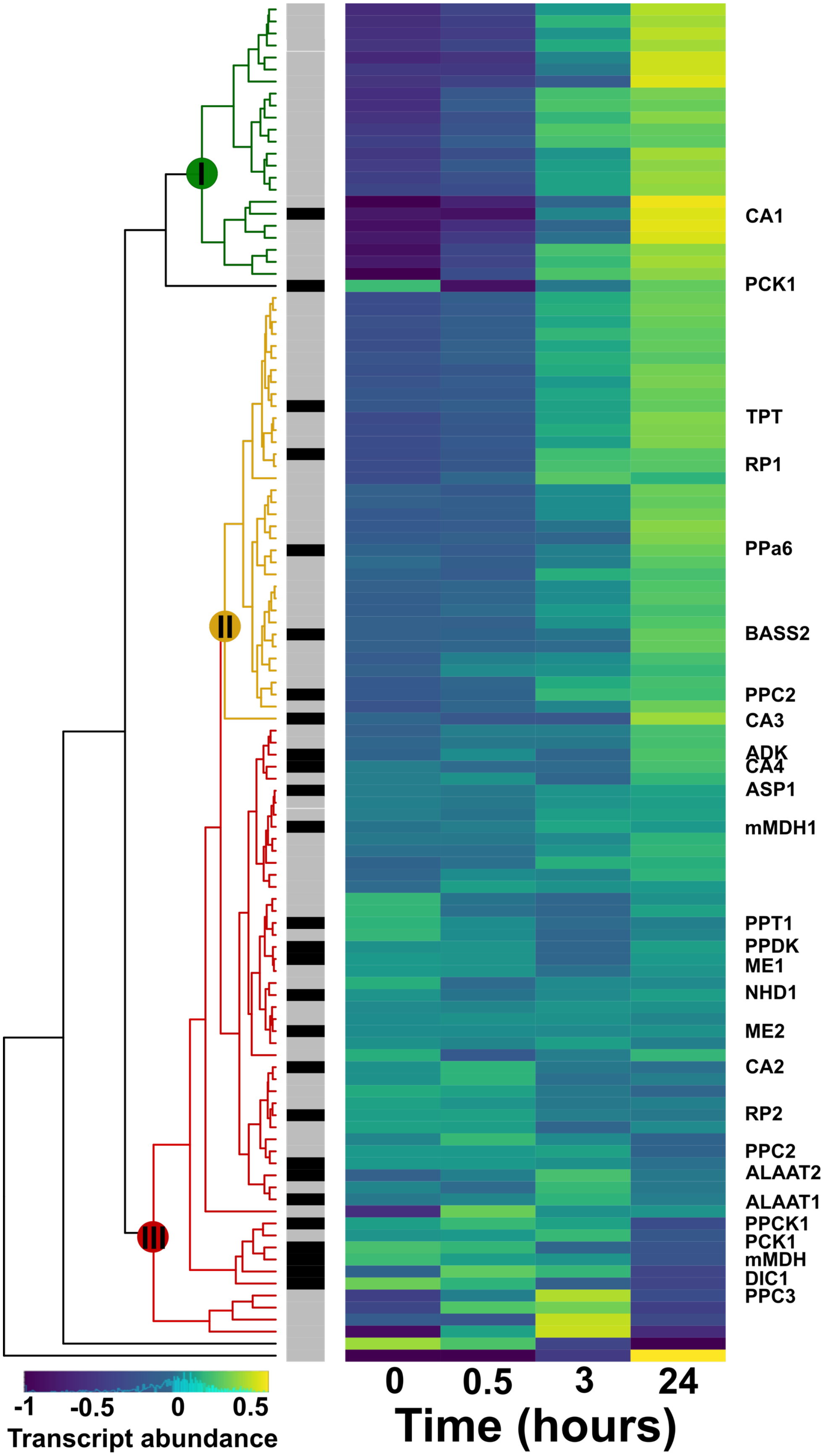


**Supplemental Figure 3:** Expression patterns of photosynthesis genes (grey sidebar) and $C_4$ genes (black sidebar) of *A. thaliana* during de-etiolation. Heatmap illustrating gene expression with each gene being represented by a row, and data centred around the row mean. Dendrograms (red, yellow and green) highlight distinct expression clusters representing strong, moderate or no induction respectively.

### Supplemental Figure 4

#### DNaseI-SEQ

##### DNaseI-SEQ Reads

| Sample | NMAP (Number of mapped reads, with a MAPQ score >42) | Signal Portion of Tags Fraction of tags that fall into a DHS from a sample of 5 million reads (SPOT > 0.25) |
|---|---|---|
| 0H | 240347172 | 0.68 |
| 0.5H | 207680388 | 0.32 |
| 2H | 211382604 | 0.28 |
| 4H | 240288496 | 0.38 |
| 24H | 245832318 | 0.36 |

##### DNase Hypersensitive Site (DHSs)

| Sample | No. of DHS | Mean DHS (in bp) |
|---|---|---|
| 0H | 249956 | 648.5 |
| 0.5H | 136187 | 532.9 |
| 2H | 96478 | 571.3 |
| 4H | 166426 | 628.1 |
| 24H | 145970 | 674.9 |

Deproteinized genomic DNA extraction from different de-etiolating time course samples

↓

Bias is calculated for each time point by dividing frequency of cuts for each 6-mer by background frequency

↓

| 6-mer | 6-mer freq. in genome | DNaseI cut freq. | Cutting bias |
|---|---|---|---|
| AAAAAA | 0.05 | 0.01 | 2 |
| AAAAAC | 0.06 | 0.05 | 0.83 |
| AAAAAG | 0.05 | 0.04 | 2 |
| - | - | - | - |
| - | - | - | - |
| TTTTTT | n | n | n |

↓

FootPrintMixture.R incorporates bias file to correct DGFs for DNase-I cutting bias

↓

DGF with Footprint Likelihood Ratios (FLR) <0 are filtered out

##### Digital Genomic Foot-printing after filtering

| Sample | DGF | DGF removed | %DGF removed |
|---|---|---|---|
| 0h | 151584 | 17634 | 11.6 |
| 0.5h | 20568 | 7712 | 37.5 |
| 2h | 25563 | 8868 | 34.7 |
| 4h | 37896 | 6480 | 17.0 |
| 24h | 131874 | 26699 | 20.3 |

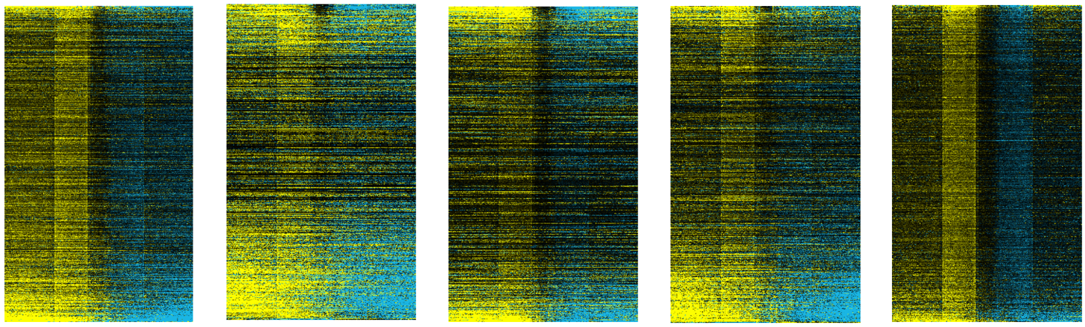

**Sample** **Dark** **0.5h** **2h** **4h** **24h**

**Supplemental Figure 4:** Pipeline for DNaseI-SEQ data processing. On the top left-hand side, pooled reads went through quality control before DHS identification using MACS2 peakcalling. The DHSs, representing accessible chromatin regions, are then searched for DGF using the pyDNase package. These DGF are prone to distorting effects due to DNaseI bias in gDNA digestion. On the top right, the pipeline for identifying this bias is shown which includes the DNaseI digestion of deproteinised ("naked") gDNA. This generates 6-mer frequencies at each cut site which is used as input for the FootPrintMixture.R tool which scores the Footprint Likelihood Ratio (FLR) of each DGF (likelihood of being a true positive). DGF with FLR < 0 were removed leaving a final set of DGF which were used for analysis. Heatmap of cut patterns are shown centred around each DGF.

### Supplemental Figure 5

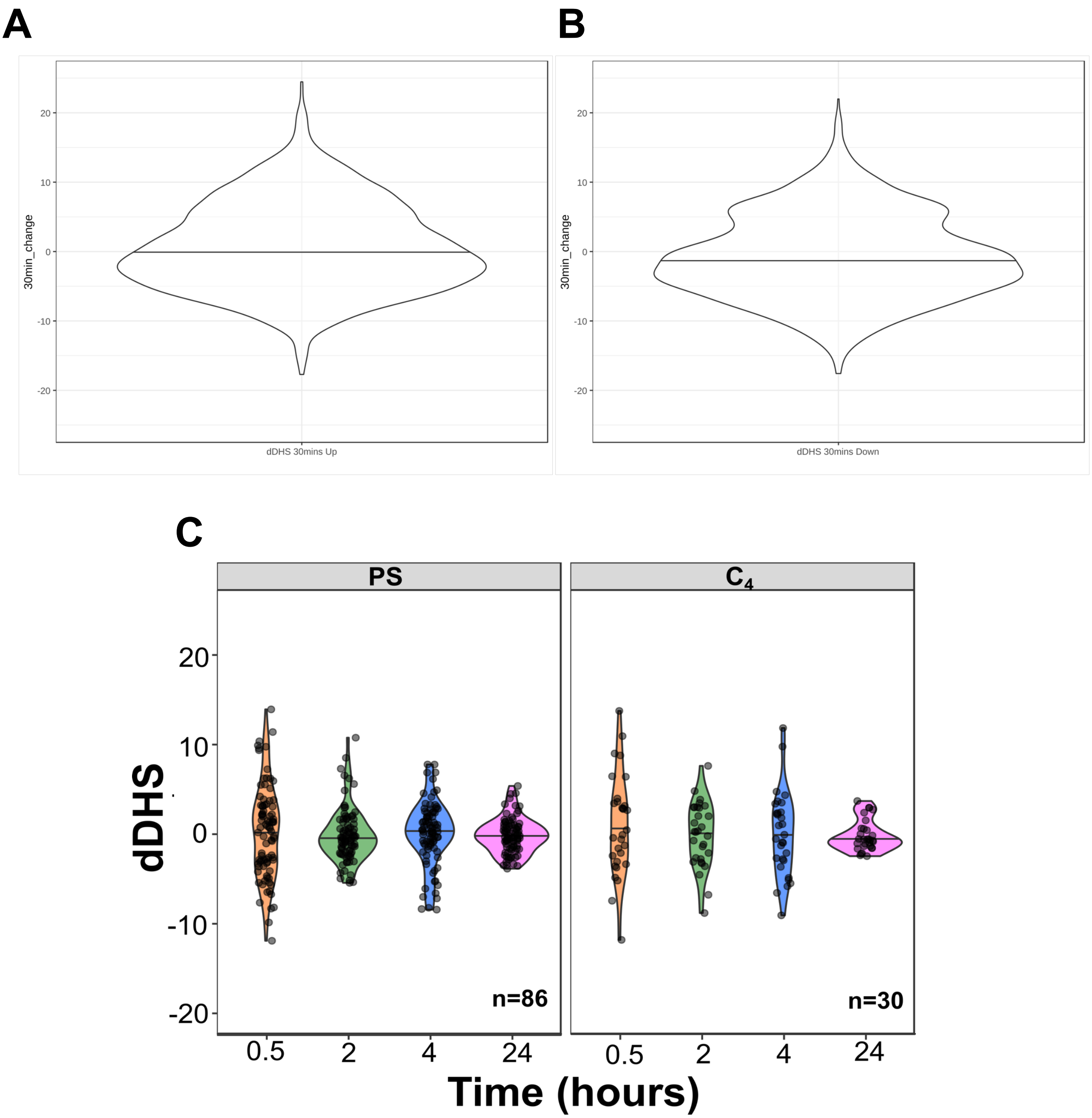


**Supplemental Figure 5**: Violin plots showing the distributions of dDHS scores for DHS overlapping with differentially expressed genes at 0.5 hours with both up-regulated (A) and down-regulated shown (B). Mean values are shown as line. Positive dDHS scores represent an increase in DHS accessibility and negative values represent the opposite. No clear association is observed with upregulated $C_4$ genes and positive dDHS values nor down-regulated genes with negative dDHS values.Violin plots depicting changes in DHS accessibility (dDHS) associated with photosynthesis genes, $C_4$ photosynthesis genes. Changes are relative to the previous timepoint, n values are for the number of DHS regions quantified.

### Supplemental Figure 6

| | ID | Targets | Features | Time-Points |
|---|---|---|---|---|
| **LOB/AS2** | Cg00747.1 | TPT | [CDS], [CDS] | [24], [0, 24] |
| | Cg00885.1 | TPT | [CDS], [CDS] | [0,2], [0,2] |
| | Cg03768.1 | CA1/2 | [Intergenic], [CDS/intron] | [4], [4, 24] |
| | Cg04272.1 | BASS2 | [Intergenic] | [0] |
| | Cg18367.1 | NAD-ME1 | [CDS] | [0,2,4,24] |
| | Cg24890.1 | PPT1 | [CDS] | [24] |
| | Cg25455.1 | PPCK1 | [Intergenic], [Intergenic] | [0], [24] |
| **Homeodomain** | Cg01471.1 | CA3/4 | [Intron] | [4] |
| | Cg04272.1 | BASS2 | [Intron] | [24] |
| | Cg05145.1 | PPDK | [CDS/Intron] | [2] |
| | Cg16610.1 | DIC1 | [Prom] | [2] |
| | Cg16884.1 | PPa6 | [Prom], [Prom], [Prom], [Prom] | [2], [0,2,4,24], [0,24],[0,24] |
| | Cg24087.1 | NHD1 | [Intergenic], [Prom] | [0], [0] |
| | Cg25455.1 | PPCK1 | [Intergenic] | [24] |
| **I-box** | Cg16884.1 | PPa6 | [Prom] | [0,2,4,24] |
| | Cg18367.1 | NAD-ME1 | [Intergenic] | [0] |
| | Cg05649.1 | ALAAT | [Intergenic] | [0] |
| **EE** | Cg00885.1 | TPT | [Intron] | [0] |
| | Cg01471.1 | CA3/4 | [Intron] | [4] |
| | Cg05145.1 | PPDK | [CDS] | [2,24] |
| | Cg16884.1 | PPa6 | [Prom], [Prom], [Prom] | [2], [0,2,4,24], [0,24] |
| | Cg18367.1 | NAD-ME1 | [Intergenic] | [0] |
| | Cg25455.1 | PPCK1 | [Intergenic], [Intergenic] | [24], [0] |
| **GT-box** | Cg05087.1 | NAD-ME2 | [CDS] | [24] |
| **G-box** | Cg01471.1 | CA3/4 | [Intergenic], [Prom] | [0], [24] |
| | Cg03768.1 | CA1/2 | [CDS] | [30] |
| | Cg04272.1 | BASS2 | [Intergenic] | [0] |
| | Cg05145.1 | PPDK | [CDS] | [4] |
| | Cg16610.1 | DIC1 | [CDS] | [0] |
| | Cg16884.1 | PPa6 | [Prom] | [4] |
| **E-box** | Cg00885.1 | TPT | [Prom] | [0.5] |
| | Cg01471.1 | CA3/4 | [Prom] | [24] |
| | Cg03768.1 | CA1/2 | [CDS] | [0,4] |
| | Cg05649.1 | ALAAT | [Intergenic] | [4] |
| | Cg18367.1 | NAD-ME1 | [CDS] | [4] |
| **C2C2-GATA-box** | Cg05649.1 | ALAAT | [Intergenic] | [0] |

**Supplemental Figure 6**: DGF binding in different $C_4$ cycle genes during de-etiolation time course in *G. gynandra*.
